# A deep hierarchy of predictions enables assignment of semantic roles in online speech comprehension

**DOI:** 10.1101/2022.04.01.486694

**Authors:** Yaqing Su, Lucy J. MacGregor, Itsaso Olasagasti, Anne-Lise Giraud

## Abstract

Understanding speech requires mapping fleeting and often ambiguous soundwaves to meaning. While humans are known to exploit their capacity to contextualize to facilitate this process, how internal knowledge is deployed on-line remains an open question. Here, we present a model that extracts multiple levels of information from continuous speech online. The model applies linguistic and nonlinguistic knowledge to speech processing, by periodically generating top-down predictions and incorporating bottom-up incoming evidence in a nested temporal hierarchy. We show that a nonlinguistic context level provides semantic predictions informed by sensory inputs, which are crucial for disambiguating among multiple meanings of the same word. The explicit knowledge hierarchy of the model enables a more holistic account of the neurophysiological responses to speech compared to using lexical predictions generated by a neural-network language model (GPT-2). We also show that hierarchical predictions reduce peripheral processing via minimizing uncertainty and prediction error. With this proof-of-concept model we demonstrate that the deployment of hierarchical predictions is a possible strategy for the brain to dynamically utilize structured knowledge and make sense of the speech input.

## Introduction

Understanding speech is a non-trivial feat. To extract information from ever-changing acoustic signals, our brains must simultaneously “compress and recode linguistic input as rapidly as possible” for multiple representation levels (1), while also keeping information in memory as we incrementally build up the meaning of an utterance (2). No computational framework to date has captured the transformation from continuous acoustic signal to abstract meaning: most speech processing models focus on either the lower-level recognition from acoustic to lexicon (3–7), or the higher-level linguistic manipulations without taking into account the constraint of elapsing time (8–13).

In addition to the challenge of fleeting time, speech signals are often ambiguous. However, humans exhibit extraordinary flexibility in making sense of ambiguous speech. We constantly make inferences based on our internal linguistic (e.g. syllabic composition of a word) and nonlinguistic prior knowledge (e.g. speaker identity, semantic context) that are learned from our personal experience. The influence of internal (prior) knowledge on speech perception takes place at all processing levels, e.g. filling the gap of possibly obscured acoustic details (14–16), or interpreting a sentence containing semantically ambiguous words (17, 18). Understanding how internal knowledge is integrated with external input on the fly is key to deciphering speech processing in the brain, and explaining the flexibility in human speech comprehension.

With the development of powerful neural networks (19–21), it is now possible for a model to implicitly learn structured linguistic knowledge from an immense amount of written text, and apply such knowledge in language tasks such as coherent text generation. Despite their remarkable achievements in specific language tasks, these models are very resource-demanding and often make egregious errors showing that their performance is not rooted in human-like understanding of the language content (22, 23). Especially if trained and evaluated on tasks involving predicting the next input (20, 21), e.g. a word, it is virtually impossible for such models to capture the abstract processing necessary for human language comprehension extending beyond linguistic forms and across cognitive domains (24, 25). A key aspect of speech understanding consists of applying structured internal knowledge to extract relevant information from the input signal. How and what internal knowledge is deployed depends on the listener’s behavioral goal, which can range from “understanding the message intended by the speaker” during a conversation to simply “predicting the next word” during an experimental task. A language model exploiting built-in linguistic as well as nonlinguistic knowledge, and driven by a behavioral goal, may hence be more powerful and polyvalent than one based on recognition and short-range prediction.

Here, we propose a computational framework in which the use of linguistic and nonlinguistic contextual knowledge allows the incremental extraction of multi-level information from the continuous speech signal. The model achieves single-sentence understanding by assigning appropriate values to semantic roles and making reasonable judgements about the nonlinguistic context in which the sentence takes place. Such a process relies on a probabilistic generative model that uses its linguistic and nonlinguistic knowledge to incrementally compose sentences. The generative model has a top context level that determines 2^nd^-level semantic roles, which are translated into a 3^rd^-level lemma sequences via linearized syntax rules. Each lemma produces a sequence of continuous, bottom-level spectro-temporal patterns via two intermediate hierarchies, integrating a syllable model (26) that was adapted from a biophysically plausible model of birdsong recognition (27, 28). Importantly, context and semantic states are maintained throughout the sentence but interact at the lemma rate, allowing the inverse model to modify previous estimates of these states with incoming evidence. During model inversion, top-down and bottom-up messages alternate at timescales of corresponding hierarchies, providing a possible solution to the “now-or-never” bottleneck (1) that is also consistent with the predictive coding hypothesis of perception (29–31).

With a small scope of knowledge adapted from stimuli in MacGregor et al. (32), the model can extract contexts and semantic roles from ongoing speech signals and resolve semantic ambiguity using new information; its beliefs about context and semantic roles, in turn, dynamically influence message passing in lower levels. The linguistically informed model structure allows for hierarchy-specific computational metrics that provide a more interpretable and holistic explanation of neural speech responses than using next-word prediction statistics generated by GPT-2 (20), a large-scale natural language model. In addition, we show that the prediction-update mechanism offers the flexibility to balance between amount of processing and inference accuracy through the control of weighting for bottom-up sensory cues versus top-down predictions.

This proof-of-concept model demonstrates a possible computational scheme of speech processing in the brain in which top-down prediction serves as a key computational mechanism for information exchange between hierarchies, driven by the goal of comprehension. Furthermore, correlations between model-derived metrics and neural responses may provide insights into the functional roles of various neuronal signals during speech perception.

## Results

### A deep hierarchical model of speech comprehension

We developed a model of speech processing based on the idea that the goal of the listener is to understand the message conveyed by an utterance. Appropriate understanding entails retrieving useful information from the utterance and optimally mapping it to the listener’s knowledge of the world, not restricted to linguistic representations (Fig 1A). Our model of the listener’s internal knowledge therefore consists of two parts that are both implemented as probabilistic generative models. The first part exemplifies knowledge about the world by defining events and properties constrained by specific nonlinguistic, situational contexts. For example, under the context of a tennis game, the listener knows (that the speaker knows) about special winning serves, about runs to return a ball etc. The serve or the run may be the central role in an event of winning a game, or described as having a certain property (e.g.: being surprising). Under the different context of a poker game, the listener knows some cards in the deck that can also be part of an event or entail some property. The second part of the model converts these events or properties into linguistic forms by choosing between a number of possible lemmas in an appropriate order, e.g. the special winning serve can be expressed as a single word “ace” early in the sentence, and finally into spoken utterances in the form of spectro-temporal sound patterns via a deep temporal hierarchy (Fig 1B). These two parts are hierarchically linked via semantics and syntax. The inversion of this generative “world knowledge” model fulfills the mapping from the sound patterns to abstract semantic roles and contexts by estimating the probability of every possible value (*state*) of each element (*factor*) in the knowledge hierarchy (Fig 1A), thus providing the listener with the means to understand the utterance produced by the speaker.

**Fig 1.**
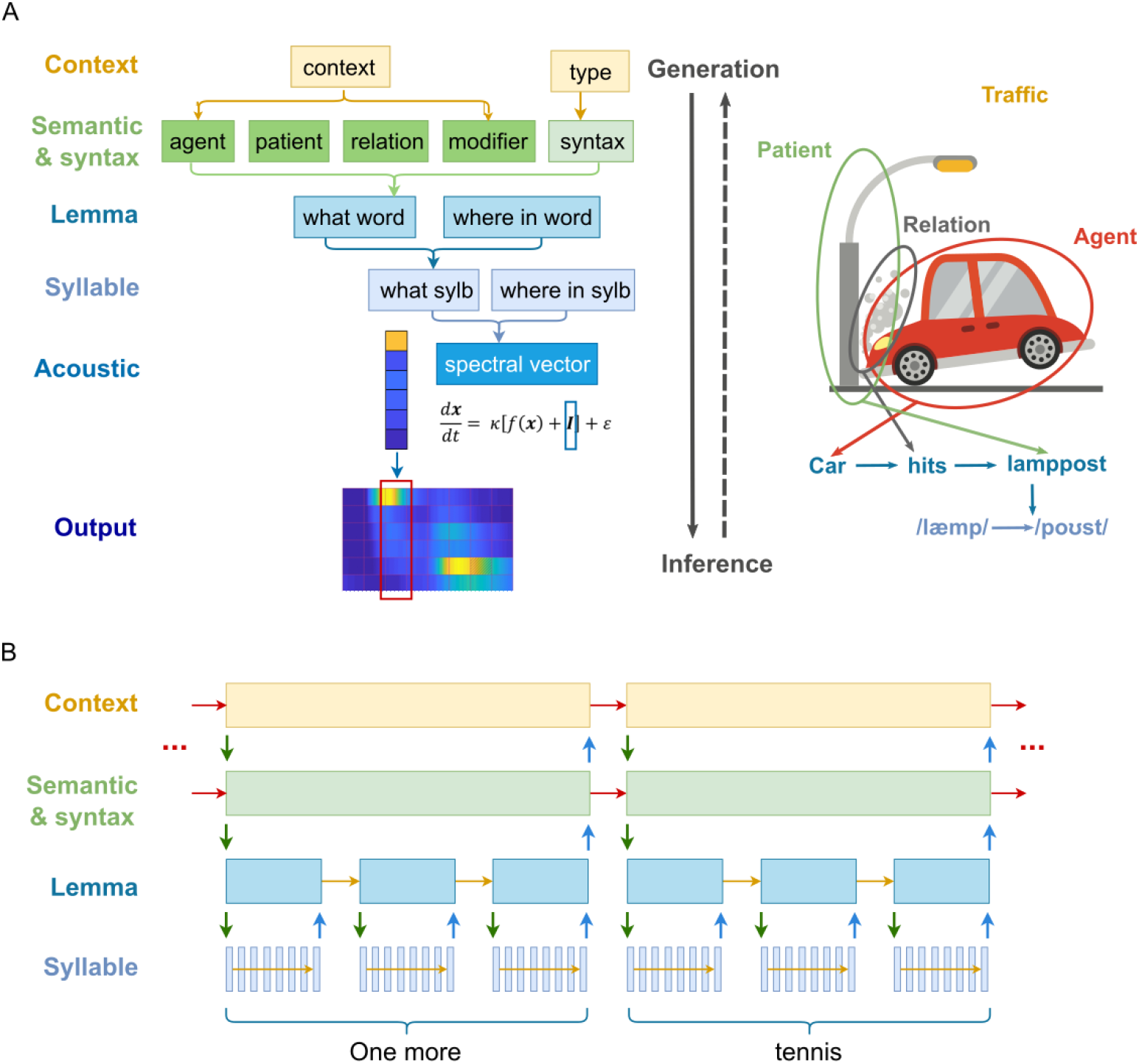
A generative model of speech and its inversion. **A. Schematic of the generative model. Left:** information conveyed in a speech signal is roughly separated into six hierarchies. To generate speech (solid downward arrow), the model first assigns values to semantic roles according to the contextual knowledge and determines a (linear) syntactic structure from the type of the message it’s expressing. Together, semantics and syntax generate an ordered sequence of lemma units. Each lemma unit generates a sequence of syllables, which in turn generates a sequence of spectral vectors. Each spectral vector unit is then deployed as a continuous acoustic signal of 25 ms. Inference corresponds to the inversion of the generative process (dashed upward arrow). The model is divided into three parts that were implemented with different algorithms (see Methods). **Right:** cartoon (www.publicdomainpictures.net) illustrating how a sequence of syllables ‘/læmp-poυst/’ (lamppost) is generated from a traffic scene context. In describing a traffic accident, the speaker tries to convey its mental image of the scene consisting of an agent (the car), a patient (the lamppost) and the relation (the action of hitting) from the agent to the patient. With English vocabulary and grammar, it chooses one lemma corresponding to each element in the accident, and outputs (speaks) these lemmas in a specific order according to the syntactic rules. Each lemma is then expressed as a specific sequence of syllables. Importantly, the same lemma can be the result of different combinations of abstract information and syntactic rules. For example, in the sentence “The ball hits the floor”, the word “hits” implies a different action than a car hitting a cyclist, whereas in “His songs are top hits” the relative position of the word implies an entity, not an action. **B. Temporal scheduling of hierarchical message passing during speech perception.** The generative model is inverted by alternating top-down prediction (prior, green downward arrows) and bottom-up update (blue upward arrows). A supraordinate level initiates a sequence of evidence accumulation in its subordinate level and receives a state update at the end of such sequence. It then makes a transition and sends an updated prediction to the subordinate level and initiates another sequence of evidence accumulation. Such a process is repeatedly performed until the end of the sentence. Note that for the lemma and lower levels, states are generated anew each time when the supraordinate level makes a transition, i.e. no horizontal arrows between sending up an update and receiving a new prior. For the top two levels, however, states are maintained throughout the sequence (red horizontal arrows) or make transitions according to a set of rules (syntax).

In all, the model includes five levels, each consisting of several factors (represented in rectangles in Fig 1A) which have multiple possible values (states) listed in Table 1 except for the *acoustic* factor, which is a real-valued vector representing the signal amplitude of six acoustic channels. Probabilistic mappings and transition probabilities between the values of the discrete factors in Table 1 are defined in Methods and Appendix. The final output of the generative model (i.e. the input to the perception model) is the continuous spectro-temporal pattern of the speech signal sampled at 1000 Hz and divided into six frequency channels (see Methods). Lengths of stimuli are fixed: each sentence consists of 4 lemmas, each lemma of 3 syllables, and each syllable of 8 spectral vectors. Every spectral vector is deployed into 25ms of time-varying continuous signal, thus each syllable effectively has a duration of 200ms (33).

**Table 1.**
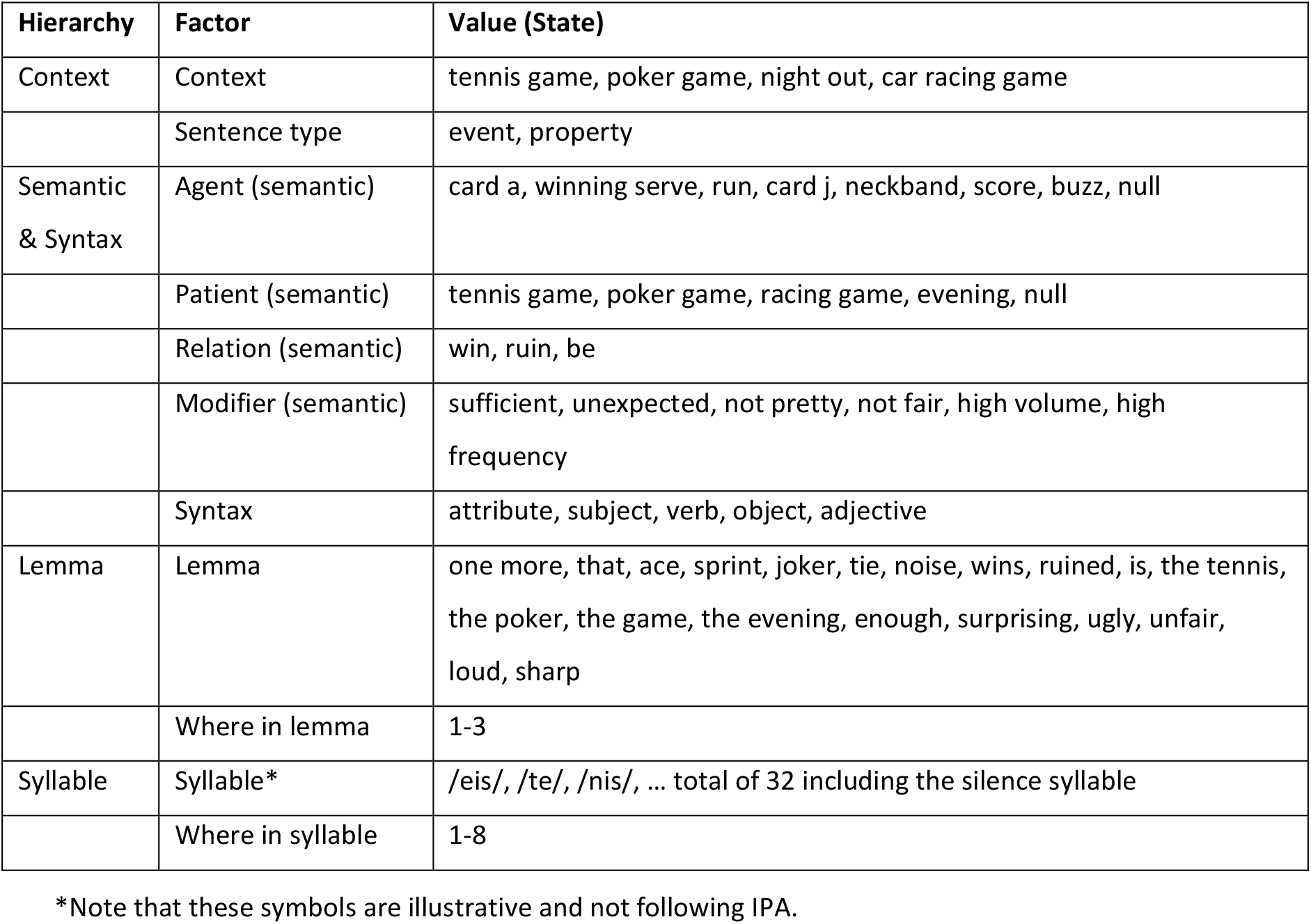
Factors and their possible values (states) in the model hierarchy.

Next, we show how this model understands simple sentences and deals with semantic ambiguity, and we demonstrate the role of top-down predictions in these processes. We assessed its performance with different sentence stimuli and parameter settings, namely by varying the perceptual bias among different contexts and the precision of the continuous module (see Methods), focusing on: 1) the probability distributions that describe the model’s beliefs (or predictions) about possible states over time, and 2) divergence and entropy measures, which summarize informational changes underlying the evolution of beliefs (see Methods). These measures do not depend on the precise fine tuning of the model parameters, and are qualitatively evaluated by whether the timing (when certain states are updated) and the outcome (what the current beliefs are about different states) of the hierarchical inference concurs with human behavior in the language domain.

Stimuli are adapted from MacGregor et al. (2020) (32) and illustrate the use of internal knowledge to disambiguate speech. All sentence stimuli in the following sections share the same structure (see Table 2 for a complete list of possible sentences):

> One more [MIDDLE WORD] wins [END WORD].

**Table 2.** All possible sentences in the model.

| Attribute | Subject | Verb | Object/Adjective | Context |
| --- | --- | --- | --- | --- |
| One more/that | ace | wins | the game/**the poker** | poker game |
|  |  |  | the game/**the tennis** | tennis game |
|  |  | is | surprising/enough | poker or tennis game |
|  | sprint | wins | the game/the tennis | tennis game |
|  |  | is | surprising/enough | tennis game |
|  | joker | wins | the game/the poker | poker game |
|  |  | is | surprising/enough | poker game |
|  | tie | ruined | the evening | night party/racing game |
|  |  |  | **the game** | racing game |
|  |  | is | **ugly** | **night party** |
|  |  |  | **unfair** | **racing game** |
|  | noise | ruined | the evening/the game | night party/racing game |
|  |  | is | loud/sharp | night party/racing game |

The MIDDLE WORD can have either one or multiple possible meanings, each meaning pointing to one context of the sentence. The END WORD either resolves the semantic ambiguity raised by the middle word or not. A disambiguating end word can also follow an unambiguous middle word without affecting its interpretation.

### The use of knowledge about the world to interpret speech

We first test how the model processes speech stimuli, with a focus on the timing of the incremental estimation process at the context and semantic levels, where “meaning” is extracted by assigning values to semantic roles.

Consider the following two sentences, A: “One more ace wins the tennis.” and B: “One more ace wins the game.” Both sentences contain the ambiguous word “ace”, which can be associated with a special serve in tennis or a special card in a poker game. The final word in the first sentence disambiguates “ace” to mean a special serve because “the tennis” can only be generated from a tennis game context, which applies to the whole sentence including the preceding “ace”. In the second sentence, however, the ambiguity remains unresolved; the game can still refer to a tennis or a poker game. In the latter case, the interpretation of the word “ace” will depend on the listener’s preference. Unless specified otherwise, we introduce a prior preference for the poker context to reflect the preference of the general population (32).

The word “ace” introduces ambiguity because it points to two possible states for *agent* (“tennis serve” or “card A”), each of which points to a separate state for *context* (“tennis game” or “poker game”, Table 2, ambiguous and disambiguating words in red). Figs 2A and 2B show the evolution of the model’s beliefs about context and semantic factors for the two sentences. The ambiguity is reflected in the posterior estimates of *agent* and *context* between the offset of “ace” and the sentence ending word, where the model assigned nonzero probabilities to “card A” and “serve” as the *agent*, and “poker game” and “tennis game” as the *context*, and near-zero probabilities for other states (Fig 2A). Probabilities for poker-relevant states were higher (darker colors) due to the contextual preference. The verb “wins” did not change the model’s estimation for the *agent* or the *context*, but clarified the sentence *type* to be “event” and the *patient* to be nonempty, again with a preference towards poker. After the model heard “the tennis” (Fig 2A), it immediately resolved its beliefs of the *agent*, the *patient* and the *context* to the opposite of its prior preference. When the sentence ended with “the game”, (Fig 2B), the model followed its preference with enhanced beliefs as a result of the entropy reduction entailed by belief updating, but not as clearly resolved as with “the tennis” (see next section).

**Fig 2.**
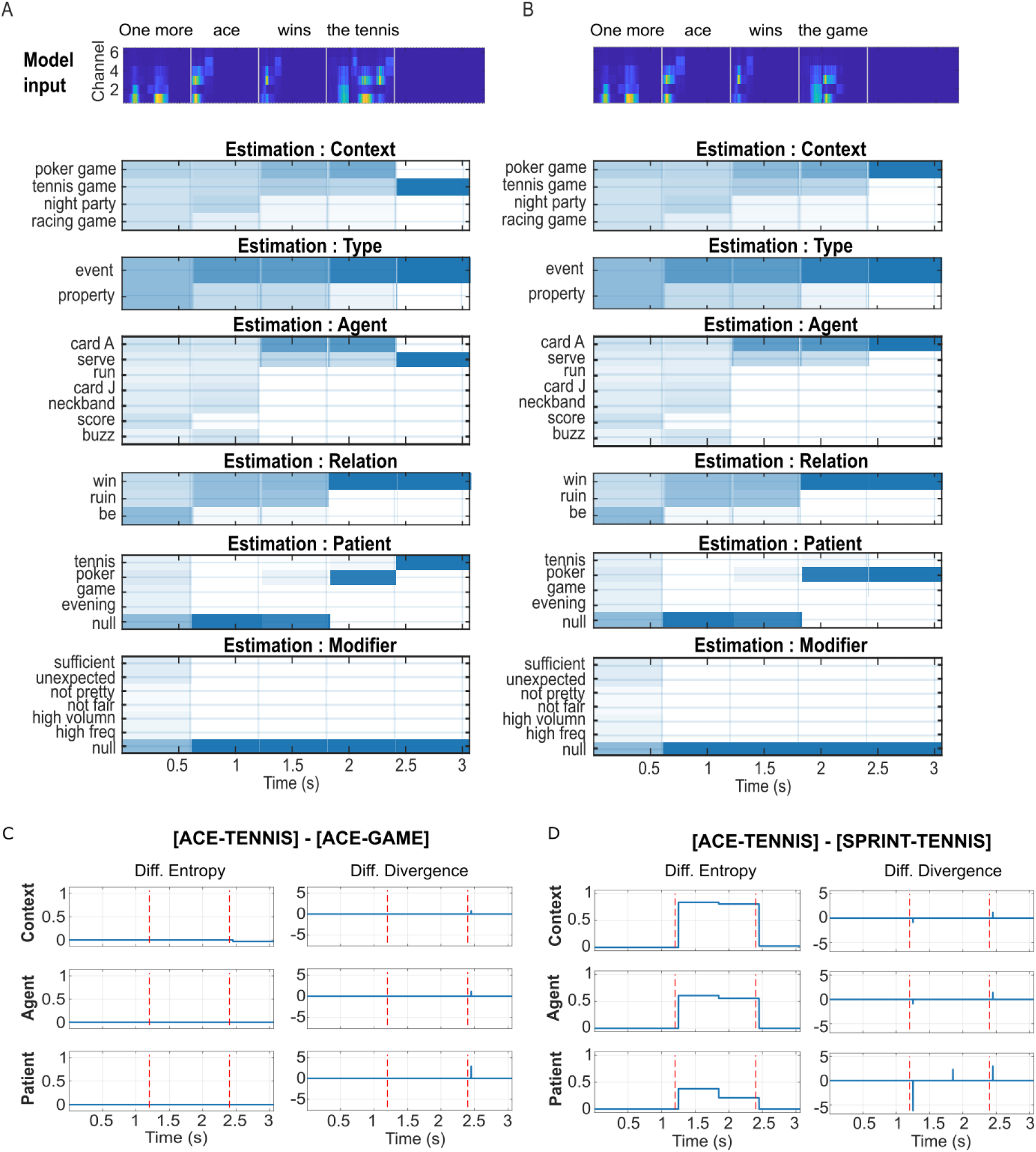
Semantic- and context-level model response to different sentence inputs. For all simulations, relative prior for context was set at the default of 1.5:1:1:1 for the four possibilities {‘poker game’, ‘tennis game’, ‘night party’, ‘racing game’}. **A. Top panel:** acoustic spectrogram of input sentence A: “one more ace wins the tennis”. Vertical grey lines mark the offset of each lemma, at which point updates were sent from the lemma level to semantic and context. **Lower panels:** estimation of posterior probabilities for the semantic (*agent, patient, relation, modifier*) and context states as the sentence unfolds. Possible values of each factor are labelled on the y axis. Blue scale blocks indicate the probability distribution for each factor, dark blue—p=1, white—p=0. The updating process is nearly instantaneous, and the main body of the n^th^ block (epoch corresponding to one lemma) is filled with the estimates after the (n-1)^th^ update. The first input “one more” was not informative. The estimated distributions were slightly changed before and after the offset of “one more” because the model still performed gradient descent to minimize free energy. After hearing “ace”, distributions for the *context* and the *agent* converged to either “poker game” or “tennis game” for *context*, and ‘card A’ or ‘serve’ for *agent*. Within these possibilities, probabilities for the poker *context* and the ‘card A’ *agent* were higher, reflecting the prior preference. Probabilities of “tennis” or “poker” as *patient* also increased*. Type, relation*, and *modifier* remain the same as in the previous epoch. After hearing ‘wins’, possibilities for *type* converged to ‘event’, and those for *relation* converged to ‘win’. Probabilities for ‘tennis’ and ‘poker’ for *patient* further increased, with a strong bias for “poker”, while the probability of a ‘null’ *patient* decreased to zero. In the last epoch, the model received a disambiguating phrase ‘the tennis’, and all factors are resolved to the correct state with a probability close to 1. **B. Acoustic input and probability estimation for the sentence “one more ace wins the game”.** The distributions are the same as in A before the last update. In the last epoch, the model receives an input, ‘the game’, that does not resolve the semantic and contextual ambiguity. As a result, distributions were further biased towards values corresponding to the ‘poker game’ context. **C. Entropy and Divergence derived from the sentence “one more ace wins the tennis” relative to the sentence “one more ace wins the game”.** The two vertical dashed lines mark the offset of the sentence middle word “ace” and the ending word, respectively. As the two sentences only differ in the ending word, both metrics differ only at sentence offset. Compared to “the game”, which does not completely resolve the ambiguity introduced by ‘ace’”, ‘the tennis’ results in lower entropy in “context” (top left panel), indicating greater certainty about the estimate. The zero differences in entropy for agent and patient indicate that the model tends to believe in its bias for these two factors. “The tennis” also gives rise to higher divergence (right panels) at sentence offset. **D. Results from the sentence “one more ace wins the tennis” relative to “one more sprint wins the tennis”.** At its offset, the ambiguous word “ace” introduces higher entropy for all three factors compared to “sprint”, reflecting greater uncertainty about the hidden states. Uncertainty dominates divergence, which is indexed by a corresponding negative difference here. At sentence offset, entropy differences between the two sentences became minimal because the model has resolved hidden states of all hierarchies. The positive difference in divergence at the offset reflects the higher surprisal for “the tennis” when it follows “ace” compared to “sprint”.

The results in Fig 2A and 2B demonstrate how prior knowledge and preferences can dynamically influence the extraction of semantic roles and contexts from the speech signal. This influence is not only reflected in the perception of semantically ambiguous words, but also in the details of message passing that give rise to its estimates. Fig 2C contrasts the inference processes between sentence [ACE-TENNIS] and sentence [ACE-GAME] in Fig 2A and 2B using their derived information metrics ([ACE-TENNIS] relative to [ACE-GAME]), focusing on the context, the agent, and the patient factors that were most relevant for the set conditions. Fig 2D compares the same metrics between sentences [ACE-TENNIS] and [SPRINT-TENNIS]. These contrasts were based on similar comparisons in the M/EEG study of MacGregor et al. (32), where the authors identified two relevant findings. First, they showed an effect of ambiguity on the magnitude of MEG sensor-space response activations shortly after the word offset (increased activation for “ace” compared to “sprint”), which could be interpreted as reflecting increased uncertainty. Secondly, they showed a (marginally significant) effect of resolving ambiguity (increase in the difference of activation between “the tennis” after “ace” vs. after “sprint” compared to between “the game” after “ace” vs. after “sprint”), which could be interpreted as reflecting increased surprisal. Respectively, these two effects were qualitatively captured by a difference in model-derived entropy (Fig 2D, left) and Kullback-Leibler (KL) divergence (Fig 2C and 2D, right) in response to the sentence contrasts. However, a difference in entropy between two conditions is often associated with a difference in divergence but in the opposite direction, with magnitudes varying across hierarchies and across factors within the same hierarchy. Thus, both semantic ambiguity and its resolution likely involve a complex combination of computational processes of different types and hierarchies. Such a complexity is in line with the finding of MacGregor et al. (32) that the two sensor-space phenomena were localized to different but overlapping sources. Further dissociation between different computation processes should involve correlating model-derived information metrics, importantly at different hierarchical levels and factors, with source-, time- and frequency-specific responses (see Discussion).

While the direction of prior preference (e.g. poker over tennis) influences both the information passing and the perceptual outcome (the state with highest posterior probability) as shown in Fig 2, the degree of prior preference also has a subtle influence on message passing during the inference process. With the same perceptual outcome, (S1 Fig A and D, either side of bias=1), the amount of information maintained between belief updates as quantified by entropy, and the magnitude information change induced by an update as quantified by the KL divergence, both vary quantitatively with the model’s prior preference (S1 Fig B-C, E-F). Thus, model-derived information metrics provide a means to relate the variability of neurophysiological responses to the perceptual preferences of individual subjects.

### Semantic prediction influences low-level message passing

The deployment of hierarchical prediction implies that high-level (*semantic, syntax* and *context*) state estimates also dynamically influence the top-down predictions (priors) as well as the bottom-up updates at lower (*lemma, syllable* and *acoustic*) levels. Figs 3A and 3B respectively show top-down priors and posterior estimates at lemma and syllable levels with the same parameters as Fig 2A. The predictions reflect both prior knowledge and the updated estimates of superordinate levels, in agreement with recent neurophysiological evidence that high-level (word) predictions constrain low-level (phoneme) predictions (34). Posterior estimations of both levels immediately converged onto the correct states after receiving the disambiguating input, for example the second syllable in the last lemma.

**Fig 3.**
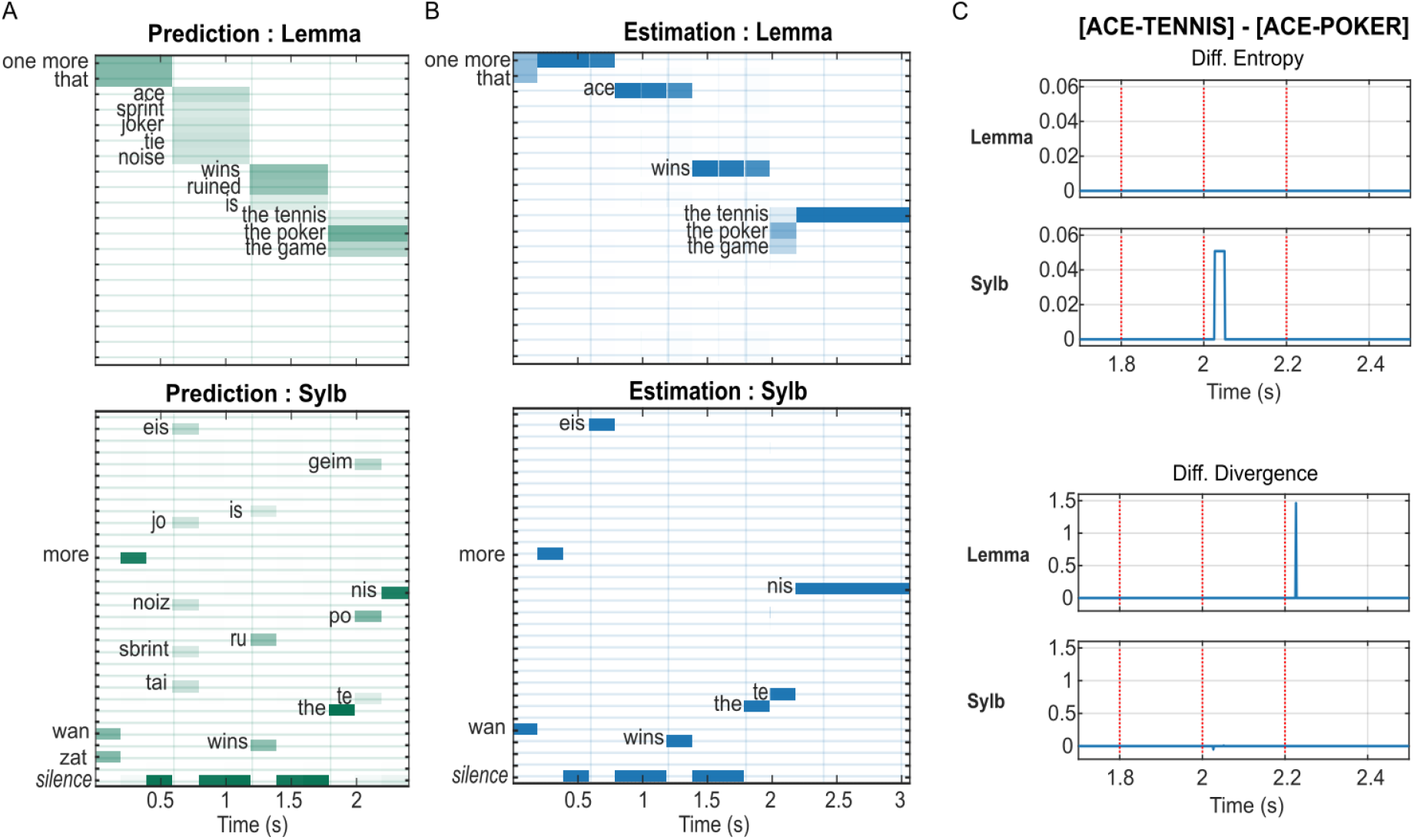
Influence of semantic state estimates on the prediction and updating of lemma and syllable states. **A. Semantic-to-lemma and lemma-to-syllable predictions (prior expectations) for the simulation in Fig 2A.** Vertical lines indicate offsets of each lemma input. In lemmas 1-3, syllable predictions (lower panel) are nearly certain after the first syllable because there was a one-to-one correspondence between the lemma and the first syllable. In lemma 4 (“the tennis”), the opposite is true because all possible lemmas start with the syllable “the”, diverging at the second syllable. Lemma predictions (top panel) depend on the current estimates at the superordinate level and the contextual bias, e.g. the prediction for the last lemma is highest for “the poker”, and lowest for “the tennis”. **B. Estimation of posterior probabilities for lemma and syllable states for the simulation in Fig 2A.** The model quickly recognizes each syllable (lower panel). The estimation for lemma states (upper panel) appears to lag for the duration of one syllable, because the lemma level receives a nearly instantaneous update at the **offset** of every syllable, and the grid between the i^th^ and (i+1)^th^ updates is filled with the estimated distribution of the i^th^ update. For example, the estimation for the first lemma started with a 1:1 prior expectation between “one more” and “that”, then converged to “one more” after hearing the first syllable “one”. The estimate was not changed until the offset of “ace”, the first syllable of the second lemma. This is only due to our graphical representation and does not affect the update from lemma to semantics. **C. Upper panels: entropy derived from sentence [ACE-TENNIS] relative to sentence [ACE-POKER] for the lemma and the syllable levels in the proximity of the final lemma.** Vertical dotted lines mark the onset of each syllable of the final lemma, either /the-te-nis/ or /the-po-ker/. A slightly higher syllable entropy after the onset of the second syllable for “the tennis” indicates the model took longer, i.e. more gradient descent steps, to converge to the less expected input /te/. **Lower panels: the difference between the divergence in response to the two sentences.** A higher lemma divergence at the onset of the third syllable (the offset of the second syllable) for the lemma “the tennis” reflects that “tennis” is less expected than “poker” due to the preference at the context level.

For the sentence input “one more ace wins the poker”, the model makes the identical semantic-to-lemma predictions as in Fig 3A (top panel), and nearly identical lemma-to-syllable predictions except for the final syllable, which was informed by the preceding syllable /po/ in “the poker” (not shown). Fig 3C shows the entropy and divergence derived from the posterior estimates of sentence [ACE-TENNIS] relative to [ACE-POKER] for the lemma and syllable level, focusing on the final lemma. Although the amplitudes of the differences are smaller than those at the semantic and the context level (Fig 2), their presence indicates that lower-level processes likely also contribute to the observed differences in neurophysiological response to semantically expected vs. unexpected speech inputs, corroborating the finding that the neural encoding of phonological and acoustic information of a word input is modulated by its semantic similarity to its preceding sentential context (34). The influence of semantic prediction on lower-level message passing can also be reflected in the processing of the same word embedded in different sentences, e.g. “the tennis” in the sentence [ACE-TENNIS] vs. [SPRINT-TENNIS] (S2 Fig). Unlike the semantic and context levels, however, the difference between “ace” and “sprint” at the acoustic and phonological levels was not reflected in the low-level message passing (S2 Fig C).

### Interpreting neural speech response requires lexical prediction and beyond

Information metrics derived from our model suggest that the sensor-space effects observed in MacGregor et al. (32) mainly reflect the message passing in semantic- and context-level processing (Fig 2), rather than in the lemma (word) level (Fig 3, S2 Fig). Meanwhile, several recent studies have successfully used word or phoneme prediction statistics derived from the output of natural language models to explain variabilities in neural response to the semantic aspects of linguistic stimuli (35–38). In doing so, the surprisal evoked by the received input given the preceding sentential context, and less often the entropy of the prediction for the upcoming input, are used directly or indirectly (in conjunction with additional regressors and regression models) as proxies of semantic knowledge to identify the neuronal dynamics underlying semantic processing. To understand the extent to which the output of a language model trained on next-word prediction can directly explain semantic- and context-level effects on neurophysiological speech responses, we reanalyzed the neurophysiological data of MacGregor et al. (32) using both explicit semantic properties as in the original study and next-word prediction statistics from GPT-2 (20) (see Methods).

We first explored whether GPT-2 predictions captured the semantic ambiguity and disambiguation in the stimuli. We adopt the terminology of MacGregor et al. (32), referring to the sentence-middle word as “Target” and the sentence-ending word as “Resolution” (Table 3). Fig 4A shows the distributions of prediction entropy after the ambiguous (blue) vs. unambiguous (orange) target word. A one-way ANOVA indicates no significant difference between entropy in the two Target word types (mean entropy: ambiguous = 4.734, unambiguous = 4.658; p=0.59). Fig 4B shows the distributions of surprisal after receiving the resolving (blue) vs. unresolving (orange) Resolution word, either following an ambiguous (left panel) or unambiguous (right panel) Target. A two-way ANOVA showed that, although the surprisal values of resolving words are significantly higher than those of unresolving words regardless of Target ambiguity (mean surprisal: resolving = 7.741, unresolving = 5.937; p<0.001), there was no difference of surprisal depending on the preceding ambiguity of the Target word (mean surprisal: prior ambiguity = 6.955, no prior ambiguity = 6.724; p = 0.74), nor on the interaction between resolution and ambiguity (p = 0.68). Thus, similar to the model’s lemma-level prediction metrics (S2 Fig C), GPT-2 entropy does not reflect the semantic ambiguity of Target words, neither does the evoked surprisal capture the long-distance interaction between Target and Resolution.

**Fig 4.**
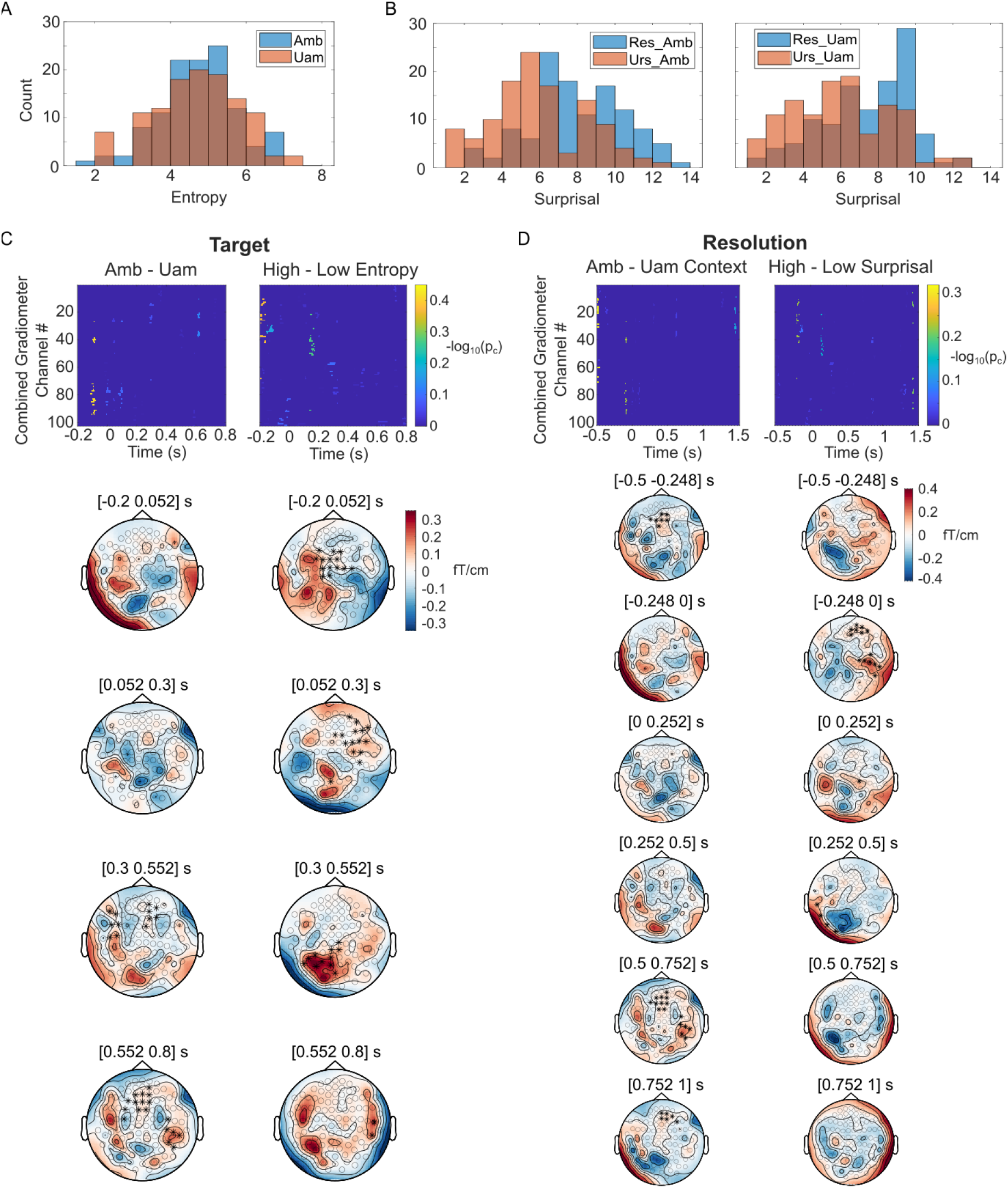
Effects of semantic properties and GPT-2 prediction statistics in MEG response to speech. **A.** Distributions of Target entropy for ambiguous and unambiguous Targets in all 58 sentences. **B.** Distributions of the surprisal values for Resolve (blue) and Unresolve (orange) Resolution words, following Ambiguous (left) or Unambiguous (right) Target words. **C. Statistical test results for the effect of semantic ambiguity (left column) and GPT-2 prediction entropy (right column) on MEG combined gradiometer data around the time of Target offset.** Top row: sensor-time maps for significance level (-log10(pc)) of sensor clusters showing the corresponding effect (see Methods for details of the calculation). Note that here both negative and positive effects are shown. Bottom rows: topological distributions of the corresponding effects averaged over four 250ms time windows spanning from −0.2 to 0.8s relative to the Target offset. Asterisks denote sensor clusters that showed a prevalent positive effect of ambiguity within the time window. **D. Statistical test results for the effect of semantic ambiguity in the preceding context (left column) and GPT-2 prediction surprisal (right column) on MEG combined gradiometer data around the time of Resolution offset.** Top row: sensor-time maps for significance level of sensor clusters. Bottom rows: Topological distributions of the corresponding effects averaged over six 250ms time windows, spanning from −0.5 to 1s relative to the Resolution offset.

**Table 3.** Example sentence input to the MEG subject and GPT-2.

| Lead in | Target | Bridge | Resolve | Unresolve |
| --- | --- | --- | --- | --- |
| The man knew that one more | ace | might be enough to win the | tennis | game |
| The woman hoped that one more | ace | might be enough to win the | tennis | game |
| The man knew that one more | sprint | might be enough to win the | tennis | game |
| The woman hoped that one more | sprint | might be enough to win the | tennis | game |

We next compared how variabilities of semantic information and GPT-2 predictions correlate with neurophysiological responses. In particular, we tested the effects of semantic properties (conceptual replication of MacGregor et al. (32)) and GPT-2 prediction statistics on the MEG response during two time windows around the Target offset and the Resolution offset. As in the original M/EEG study, we focused on combined gradiometer pairs, which demonstrated the most robust effects, and two analysis time windows around the Target offset and Resolution offset respectively.

For the Target time window, we split the MEG response into two groups according to the property of the Target word pair: 1. The GPT-2 entropy of the ambiguous Target is larger than that of its unambiguous counterpart, and 2. The GPT-2 entropy of the ambiguous Target is smaller than that of its unambiguous counterpart. S3 Fig A shows the distribution of entropy differences between ambiguous and unambiguous Target word pairs (ambiguous minus unambiguous). Ambiguous Target words with difference > 0 (i.e. in group 1, 29 pairs in total) and unambiguous Targets with difference <0 (i.e. in group 2, 29 pairs in total) contribute to the high-entropy group, and the rest contribute to the low-entropy group. Such splitting ensures that the pair of Target words in the same sentence set is always separated into two conditions, thus controlling possible confounds of the preceding sentential context. Using a data-driven algorithm (see Methods), we identified sensor-time clusters that showed a significant effect (two-tailed paired student’s t-test, p<0.05, same in the following results) of semantic ambiguity by contrasting responses to ambiguous Target vs. unambiguous Target words, (Fig. 4C, left column). We also identified clusters showing an effect of GPT-2 entropy by contrasting responses to Target words with high vs. low entropies (Fig. 4C, right column). Sensor-time statistical maps (Fig 4D, top row) as well as topographic plots over time (Fig 4D, bottom row) indicate that these two effects are likely distributed differently both in space and time. The absence of a significant correlation (Pearson’s correlation r=-0.04, p=0.66) between the sensor-wise effect sizes of the two contrasts (S4 Fig A) also suggests that semantic ambiguity and GPT-2 prediction entropy may account for different spatial aspects of the MEG responses. Interestingly, the positive effect of GPT-2 entropy arose before the word offset, whereas the positive effect of semantic ambiguity was only apparent after the word offset (Fig 4C, top row).

For the Resolution timepoint, responses to only the Resolve sentence ending were split into two groups in a similar fashion as for Target: 1. The GPT-2 surprisal following the ambiguous Target was larger than the same word following the unambiguous Target, and 2. The GPT-2 surprisal following the ambiguous Target was smaller than the same word following the unambiguous Target. Thirty-six out of the 58 sentences were labeled as being in group 1, 22 in group 2 (S3 Fig B). The contrast between Resolution words following ambiguous vs. unambiguous Target words revealed an effect of ambiguity of the previous context distributed among right temporal-parietal and midfrontal areas spanning several time windows before and after the word offset (Fig. 4D, left column). The contrast between Resolution words with high vs. low surprisal revealed an effect of GPT-2 prediction surprisal with a different spatial distribution, and restricted to −250 to 250ms (Fig. 4D, right column). Similar to the Target effects, the effect sizes of ambiguity and surprisal at Resolution offset were not correlated (r=0.001, p=0.99, S4 Fig B) across sensor locations.

These results demonstrate that both GPT-2 word-prediction statistics and high-level semantic properties contribute to the variability in neural speech responses, but their effects exhibit different spatio-temporal distributions. Given that predictions from the GPT-2 output cannot directly capture the semantic properties we investigate here (Fig 4A, B), the approach of interpreting the neural response to speech (and more generally language) solely based on such predictions learned from word sequence statistics overlooks important aspects of the dynamics underlying higher-level language processing. Our model, on the other hand, explicitly depicts multiple levels of linguistic and nonlinguistic processes under the same computational principles. Thus, it points to a more interpretable and holistic approach to characterizing the functional network underlying speech comprehension. A quantitative mapping between model and neural responses requires a nontrivial expansion of the model and is beyond the scope of the current study (see Discussion).

### Top-down prediction reduces processing effort

The model works by iteratively calculating the discrepancy between top-down predictions (expectation of the input) and bottom-up input at each hierarchical level, and using such a discrepancy to modify the state estimates of superordinate levels. This does not mean the model needs to make the best prediction for the next input as in Fig 3A: hierarchical predictions are a necessary computational mechanism in relaying information for making better inferences, even if the actual input deviates from the predicted one. To examine how the prediction content may influence the inference process, we ran the model using the same input as Fig 2A and 3B, “one more ace wins the tennis”, but simulating the extreme case of uninformative (uniform distribution across all possibilities) top-down predictions. We found that the predictive content influenced both the model time course and final estimate.

Compared to the condition of informative top-down predictions (Fig 3B), when top-down predictions were uninformative, the model still made correct inferences about every input, but with a slight delay for syllables (Fig 5A). Fig 5B contrasts the entropy and cumulative divergence during the inference process between the two conditions. Unsurprisingly, informative predictions lead to reduced entropy (maintenance of possible items) and divergence (magnitude of updates after the integration of new evidence), both contributing to fewer steps of gradient descent at each point of belief updating, hence less computation effort in terms of processing time and energy cost (39).

**Fig 5.**
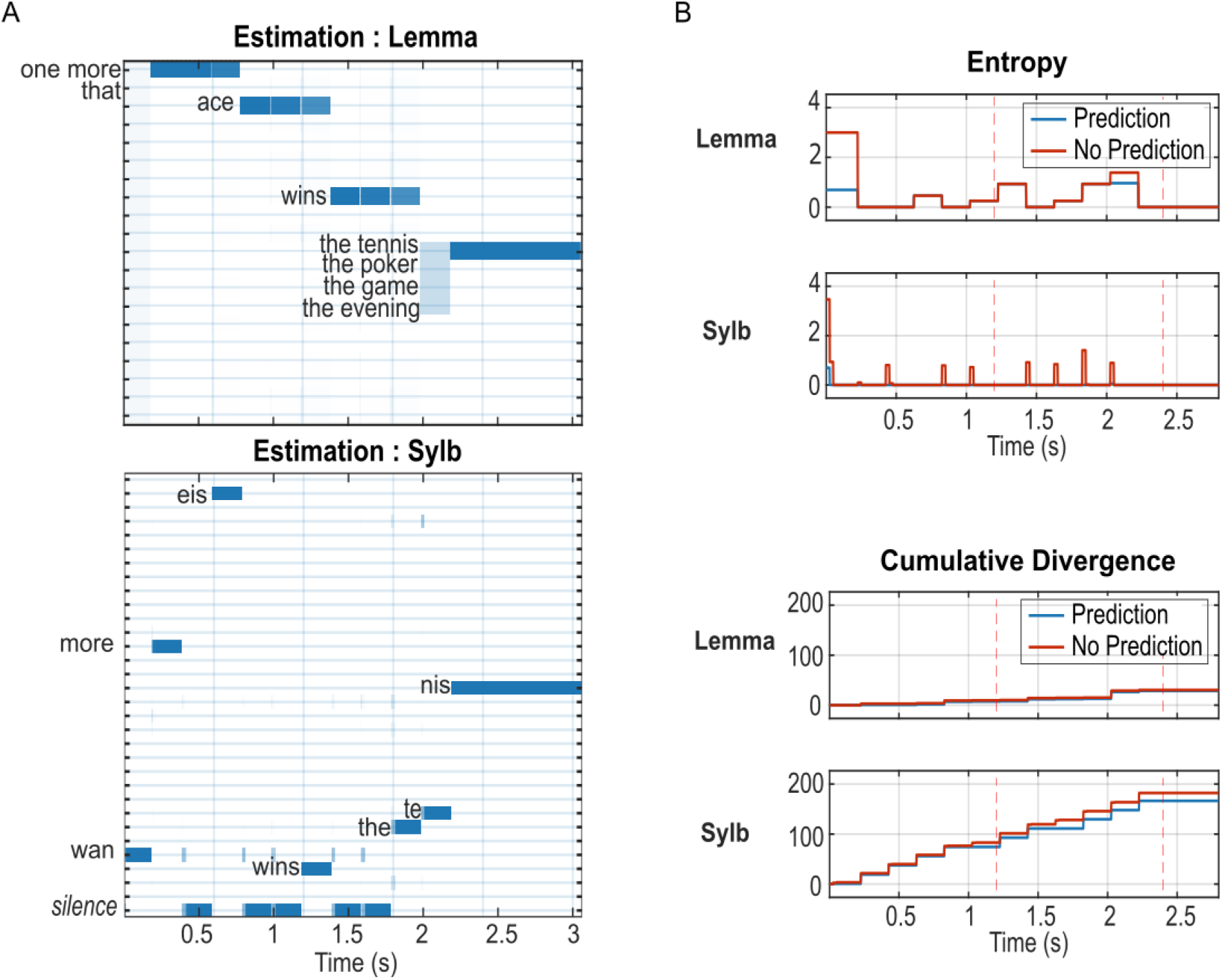
Influence of top-down predictions on syllable and lemma inference under high peripheral precision. All results are simulated with the sentence “One more ace wins the tennis”. With uninformative predictions, model responses at the semantic and context levels are nearly identical to Fig 2A because the model reached the same, almost-certain lemma estimates at the time of semantic updating (at each lemma offset). Therefore we omit the higher-level results here and in Fig 6. **A. Estimation of posterior probabilities when top-down predictions are set to uniform distributions for all possible states.** Compared to Fig 4B, there is a slight delay for the convergence of every syllable indicated by the small vertical bars, each corresponding to one spectral vector, in more than one possible state. The inference for lemma states is not significantly changed: once the model is certain about the first (or the second in the case of the last lemma) syllable, it can quickly converge to the correct lemma using its internal knowledge. **B. Upper panels: entropy calculated from lemma and syllable states.** With uninformative top-down prediction (red), the entropy of syllable states was raised for a short duration (~1-2 spectral vectors) more often than with informative (blue) prediction (eight times throughout the sentence versus once at the sentence onset). The difference is less obvious at the lemma level except during the very first syllable and the /the/ syllable in the last lemma. **Lower panels: cumulative KL divergence for the two factors.** Overall, the cumulative divergence is smaller when informative prediction is available (blue).

So far, we have simulated the model with the ideal scenario of arbitrarily high precisions (see Methods) at the continuous level. In general, a high precision implies that fine details of the input are utilized to evaluate the mapping between the input signal and the generative model, analogous to a perfect periphery that preserves the best possible spectro-temporal information from the acoustic input. It has been suggested that top-down predictions may be especially important under challenging situations, e.g. impaired auditory periphery (40). We tested the model with a broad range of precisions to assess how precision affects online speech processing. In particular, we lowered both the precision for the continuous state as well as for comparing the input with predicted activity in the six frequency channels (see Methods), which is analogous to lesioning the local computation supported by lateral connections and the cross-level information carried by bottom-up connections, respectively (28, 41). Within a considerable degree of degradation, the model performance is qualitatively the same as the intact model, in that it correctly infers the states of all factors, but a strong difference arises in the time it takes to converge, especially in the case of uninformative top-down predictions (S5 Fig A and B, precision=exp(6) vs. exp(16) in the intact condition). Fig 6 shows the comparison of informative vs. uninformative predictions similar to Fig 5, but with much lower peripheral precisions (exp(0)). Syllable identification was delayed in both cases when compared to their intact-periphery counterparts (Fig 6A vs. 4B, 6B vs. 5A), and the delay was more pronounced with uninformative predictions. This dramatic delay with uninformative prediction is accompanied by higher entropy (Fig 6C, upper panels) as well as divergence (Fig 6C, lower panels). However, an increase in effort during syllable recognition may be important to avoid inaccurate recognition: in Fig 6A, although the model saved processing time by relying on its prior knowledge, it did so at the cost of incorrectly identifying the final lemma as “the poker”. The tradeoff between processing and accuracy has been well-documented in the decision-making literature (42) and neuroeconomics (43), which reveals that humans flexibly adapt their strategy in challenging scenarios where high accuracy and low effort cannot be achieved simultaneously. Our results suggest that such tradeoff can be manipulated via adjusting one’s reliance on top-down prediction vs. bottom-up sensory information, an ability widely involved in perceptual processes including inferencing others’ intention (44) and likely lacking in certain neuropsychological disorders such as those inducing hallucinations (low sensory precision but high prediction precision) and autism spectral disorder (extraordinarily high sensory precision) (45). Nevertheless, the effort-accuracy tradeoff is also limited by the capacity of the sensory periphery: at extremely low precisions, the model’s syllable recognition breaks down without the guidance of informative top-down prediction (S5 Fig C and D, precision = exp(−4)).

**Fig 6.**
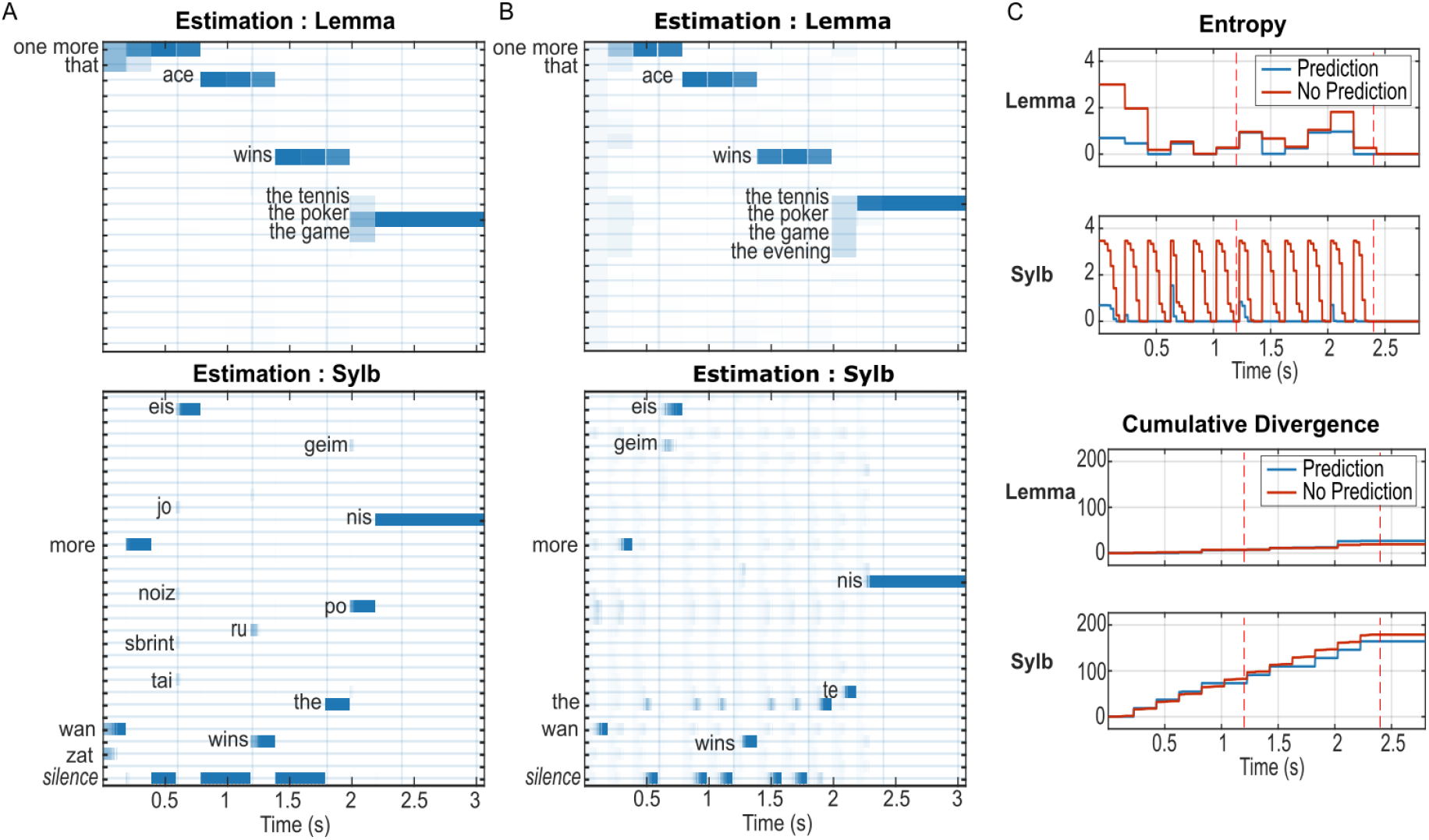
Influence of prediction with lowered peripheral precision. The input sentence, as in Fig 4B and 5A was “One more ace wins the tennis”. Precision was set to p=exp(0), whereas in the intact model (Fig 4B and 5A) p=exp(16). A and B: state estimation with and without informative prior. C and D: entropy and divergence in the two conditions. **A.** With informative prediction, the result is similar to that in Fig. 2A, except that 1) for the last lemma input, the model relied on the prediction, biased towards “the poker”, and made the wrong inference, and 2) for the starting syllable in each lemma, the model took several spectral vectors to converge as indicated by the colored bars. **B.** Without prediction, the model took longer to infer each syllable compared to A, but the inference was correct. **C. Upper panels: entropy with informative (blue) or uninformative (red) top-down prediction for lemma and syllable estimates.** Without informative prediction, the uncertainty increased at the onset of every syllable instead of only for the syllable with multiple possible candidates (e.g. the syllable after “the” in the last lemma), and also reached higher magnitude as well as longer duration compared to the informative condition. **Lower panels: cumulative divergence in the two conditions.** The divergence for syllable states was lower with informative prediction, but not for lemma. However, the summed divergence of the two levels is slightly higher with uninformative prediction.

Overall, the model demonstrates that hierarchical prediction, whether highly informative about the next input or not, can serve as a key computational mechanism for robustly extracting structured information from ongoing speech, and that informative predictions are desirable when processing effort needs to be minimized, and in time constrained situations (e.g. turn-taking). With an impaired periphery, greater effort is required to obtain accurate perception.

## Discussion

The idea that our brains adaptively entertain internal models and that this facilitates language comprehension underlies much current research in speech (language) perception. Nevertheless, how internal knowledge is deployed in time, in relation to the timing of continuous speech unfolding, is an open question, and may be key to achieve the form-meaning distinction in neural-network language models (23, 24). Here, we attempt to establish a foundational framework that dynamically exploits general knowledge in speech comprehension to bridge this gap. We implement the listener’s internal knowledge as a probabilistic generative model that consists of a non-linguistic general knowledge (cognitive) model and multiple temporally organized hierarchies encoding linguistic and acoustic knowledge. Speech perception, modeled as the inversion of this generative model, involves interleaved top-down and bottom-up message passing in solving the computational challenge of extracting meaning from ongoing, continuous speech. We show that the model makes plausible inference of hierarchical information from semantically ambiguous speech stimuli and demonstrate the influence of prior knowledge on the inference process, which is reflected in the neural response to speech stimuli but not in next-word prediction statistics of a deep neural-network language model (GPT-2) (20). We also show that hierarchical predictions can be exploited to reduce processing effort. The model tries to mimic human language comprehension by jointly implementing incrementality and prediction (46), and could potentially be expanded towards a comprehensive model of natural language *understanding*, and guide the interpretation of neurophysiological phenomena in realistic listening scenarios.

### Language comprehension as semantic role assignment

Although we emphasize that speech (language) comprehension is driven by high-level behavioral goals, to achieve comprehension the appropriate assignment of semantic roles is crucial for (re)constructing the message conveyed in the utterance, e.g. the “mental image” in Fig 1A. Semantic roles can be viewed as an interface between linguistic and nonlinguistic representations, the latter being a fundamental, domain-general format of our internal abstraction of the world (24, 25) that is shown to both behaviorally and neurophysiologically influence language comprehension (47, 48). The process of semantic role assignment is central in psycholinguistic process theories (46, 49–51), yet seldom reflected explicitly in existing computational models of language. A major challenge for modeling semantic role assignment during language processing is in combining meaning extraction with compositionality: words that carry semantic contents are presented in an order dictated by compositional rules, thus the extraction of persisting meanings must take place dynamically alongside the decomposition. These two aspects have only been addressed separately in some existing models, e.g. topic models (9, 52) fulfill (lexical) semantic processing but ignore the word order. On the other hand, the Discovery of Relation by Analogy model (11, 53) learns the time-based binding rules that decompose words and phrases into hierarchical structures, but does not have explicit representations of semantic knowledge.

A recent model of linguistic communication (12) did incorporate abstract nonlinguistic (geometric) knowledge and compositionality, but lacked the incremental nature of the meaning-building process in humans (2). The generative model encoded several templates of complete sentences and a set of geometric properties. By applying nonlinguistic knowledge under the goal of resolving object properties, the model generated sentences by picking the most probable sentence format and filling specific positions with the most helpful descriptive words. The inverse model thus comprehends a word sequence by inferring the sentence format and capturing keywords at the corresponding positions. This template-matching strategy realized a form of meaning-structure conjunction. However, it constrains the model comprehender to update its estimate of the sentence at the sentence offset instead of on the fly during the sentence.

Our model achieves human-like speech (language) comprehension in that it applies syntactic rules to dynamically update values assigned to semantic roles with each incoming lemma. It does not rely on a direct representation of sentences, but incrementally builds up its understanding of an utterance through incorporating new evidence into current beliefs of semantic roles. We share this notion with the Sentence Gestalt (SG) model of language comprehension, which achieves dynamic thematic role assignment from lexical inputs using a neural network trained on linguistic stimuli produced by a probabilistic generative model (13, 54). The function of situation and thematic roles in this generative model are homologous to that of the context (situation) and semantic (thematic) factors of our model. However, while the SG model extracts thematic information from lexical input, a central feature of our model is to deploy all the hierarchies from the *online* processing of continuous speech to language comprehension. The variational Bayesian approach and the gradient-based algorithms we used here have two particular advantages. First, it allow us to explicitly model the interactions within and between meaningful computational hierarchies, and second, they can account for dynamics of neuronal activities such as local field potentials (39, 55). We therefore believe our model is better suited to our goal of explaining language processing within a potentially unifying account of neuronal message passing, rather than in terms of neural-like network activations (see next section).

The behavioral (nonlinguistic) goal of language comprehension is implemented minimally in the current model as the task of inferring a simple context (situation) level, which represents the basic “world knowledge” necessary for resolving semantic ambiguity. To implement cognitively more elaborate language tasks, the context level in the model would need to include additional elements that likely involve multiple decision hierarchies (56). Yet, while a model can include an arbitrary number of hierarchies, there is not an infinity of corresponding specialized brain regions. Computational hierarchies, especially those of higher cognitive functions that can expand to an infinite depth, are therefore likely embodied by information exchanges among a limited number of functionally specialized regions, through reciprocal interactions that can theoretically implement unlimited hierarchical structures using only two abstract chunking levels (57, 58). These information exchanges reflect the probabilistic mappings in the comprehender’s internal model, as shown in Figs 2 and 3, and play an important role for linking the model’s computational principles to neurophysiological data of speech information processing in the human brain.

### Understanding neural information transfer through divergence and entropy

Brains process internal and external information with high efficiency. Two types of information theoretic metrics have been of particular interest in establishing the connection between abstract information and biophysical signals to probe the brain’s information processing capacity: surprisal (related to, but distinct from divergence) and entropy. Efforts in associating neurophysiological responses to surprisal for next-word expectation, either based on cloze probability tests (32, 59–61) or the probabilistic distribution estimated by computational models (35–37, 62–65), largely credit Levy’s influential work on expectation-based comprehension (10). Levy proposed a formal relationship between incremental comprehension effort and the Kullback-Leibler divergence (KLD) of syntactic structure inference before and after receiving a word input W, and proved that the KLD reduced to the surprisal of W given the previous word string when conditioned on a constant extra-sentential context that constrains comprehension. Although these studies robustly found neurophysiological correlates of word surprisal, focusing on this aggregated measure without explicitly modeling probabilistic representations above the word level may not be enough to tease out the influence of high-level factors on language processing as was shown in Figs 3, 4 and S2 (62). High-level processes presumably explain conflicting findings across studies on evoked response (66) and underlying neuronal circuits (32, 36, 61) of word surprisal, because different experimental paradigms likely tap into different language processing modes, making word surprisal too coarse a measure. Here, we demonstrate the possibility to explicitly model information transduction above lexical processing and use KLD as a universal metric to quantify information transfer, in line with some predictive coding hypothesis that propose KLD to be driving the prediction error signal transmitted between cortical hierarchies (55, 67).

Regarding entropy, the measure of information in a system (68) that represents the uncertainty in linguistic stimuli, it has drawn less interest compared to surprisal metrics (32, 36, 69, 70). There is no consensus on how information is maintained between two instantaneous belief updates, and entropy may be valuable in investigating the *representation* of information in the brain. Intuitively, higher entropy implies greater effort (more possibilities to be maintained), and less precise estimates thus weaker top-down prediction influence, but it is unclear what neural activities can underpin such effects. Noninvasive whole-brain imaging may inform us when and where the effort takes place given that entropy and divergence can be properly dissociated (36), whereas the biophysical implementation, e.g. neuronal firing patterns, may only be revealed by invasive methods.

By showing that information passing across different processing levels contribute in a complementary manner to the variability of the neurophysiological response to speech (Fig 4), our model supports the neural processing of language as hierarchically organized information passing among brain areas. Both KLD and entropy, as well as bottom-up prediction errors and top-down priors that can be decomposed from KLD (71), are suitable metrics for such an investigation. Although no definitive conclusion has been drawn on the anatomical circuits involved in high-level (semantic and beyond) message passing during speech perception, a converging view is that the extraction of different hierarchical representations is distributed in networks that perform multiple subprocesses in parallel (72–75). Recent temporally and spatially resolved neuroimaging studies suggest that neural oscillations are a good candidate mechanism for timed information transmission in these subprocesses (67, 76–78). The discrete portion of our model, or in theory any model with explicit structural and timing information (11, 53), can provide a template for organizing distributed oscillatory activities into functional hierarchies through correlating latency- and frequency-specific neuronal dynamics with model-derived information metrics. In general, sensory inputs sampled by fast (gamma) oscillation are parsed into higher-level information as phase alignments of slow (theta, delta) oscillations (26, 76, 79–82), which are found to be modulated by level-specific speech information (32, 36, 61) and top-down coordination of mid-range (alpha, beta) oscillations (78, 79, 83–87). One promising avenue that exploits both model-derived computational metrics and neural oscillations to disentangle neural information transfer is via a forward model that explains the neurophysiological signal as a result of input-modulated changes in direction-specific connection strengths between specific neural sources (brain areas), i.e. effective connectivity (88, 89). Through hypothesis testing of specific brain areas and their connectivity patterns relevant for language processing, direction (top-down or bottom-up) of information transfer can be distinguished by frequency band-specific induced activities (90), and the functional hierarchy as well as the computational roles of different connections may be mapped by regressing their modulation gain with model-derived information metrics.

The proposed approach is fundamentally different from a purely data-driven one that identifies neural response patterns correlated with pooled activities from hidden layers of a neural network trained on specific tasks of next-input predictions such as in (62, 64, 65). The brain interacts with the external stimuli, whether linguistic or not, in a structured fashion that is likely reused across different domains (44, 58). Thus, a clear computational interpretation of brain activity patterns requires an explicit representation of such structures that is lacking in most neural network models.

### Future development towards natural language understanding

In this work, we provide a basic model that integrates linguistic and nonlinguistic world knowledge in speech perception. Though the current work focuses on resolving ambiguity in semantic role assignment within a reduced language and world model, the framework of a hierarchical generative model is suitable for capturing various features of human language processing. For example, additional branches can be “plugged-in” onto specific levels of the current generative model to enable multi-modal speech processing. One possible case is to generate continuous lip movement from each syllable (91, 92), in parallel with the syllable-to-acoustic generation. The inverse (comprehension) model is then equipped to deal with audiovisual speech input, and can thus potentially simulate known effects including using one modality to disambiguate the other (e.g. a high-precision visual processing to mitigate noisy auditory input), or processing conflicting bimodal inputs (e.g. relying more on the modality that has higher precision)(92). The additional branch can also be attached to the context level to generate a sequence of events, such as a car speeds up and hits a streetlight, to allow the inverse model to make inference about the shared context from both linguistic (speech) and nonlinguistic inputs.

Another important feature of language processing is learning, which is also necessary for upscaling the model to reflect the wealth of linguistic and nonlinguistic knowledge mastered by a real listener. Language learning can be conceptualized as consisting of two complementary components: 1) learning the structure of the generative model, including the possible states of different factors and syntactic rules; 2) learning the parameters of the generative model, including priors, likelihoods, and precisions, which are fixed in the current model. Although it is nontrivial to extend the current model to include either type of learning, they could be achieved within the framework of probabilistic generative models. For the first type, a plausible algorithm of statistical parameter learning of structured contextual and semantic knowledge is the one proposed for the “topic” model of semantic representation (9, 52). Griffiths et al. (9) also pointed to a possible way to integrate complex syntax and semantic generative models by replacing one component in a syntax model (93) with such a topic model. This would allow the syntax model to determine an appropriate semantic component for the current timepoint and the semantic model to generate a corresponding word, which is consistent with the way semantic and syntax factors interact in our current model. More recently, Beck and colleagues (94) showed that a formal equivalence of the topic model can be implemented via a probabilistic (neural) population code, providing a plausible path to a neural implementation of the model. The second type of learning can be viewed by updating the relevant parameters within a fixed structure learned from a structure-learning model. Such an updating algorithm has been implemented within the dynamic expectation maximization (DEM) framework that we currently use (95). To exploit the algorithm, the current generative model needs to be modified to include a relevant task and associated rewards (both external and internal), so that the model can actively adjust its parameters to optimize rewards. This way, top-down predictions can evolve from naïve (e.g. uniform prior as we simulated in Results) to specific.

Overall, this model adopts a different and complementary perspective from the rapidly developing world of large-scale natural language models (19–21) in that it puts upfront the gross biological factors that motivate language in the first place (96–99), rather than those that seek to match human performance via selected measurements in specific tasks. Recent interesting endeavors in merging these two perspectives focus on adding more “neural features”, such as longer memory span and domain-general knowledge beyond language, to improve natural language models (24, 25). While this strategy is useful from the viewpoint of artificial language processing, it stays relatively removed from the specific biological substrates of language and hence sheds little light on how human language emerged and evolved under evolutionary pressure. Here, we propose a computational framework to address more directly these fundamental questions by explicitly including nonlinguistic components in the model architecture and using hierarchical (as opposed to aggregated) prediction as a general computational strategy. Although here we focus on a passive listener, a comprehensive model of human language understanding should also consider interactive aspects of language, i.e. language production and multi-person communication (12) where language serves as a medium to achieve shared goals (24, 100–103).

## Methods

### Model for speech comprehension

We model speech perception by inverting a generative model of speech that is able to generate semantically meaningful sentences to express possible facts about the world. Since our main goal is to illustrate the cognitive aspect of speech comprehension, we use the model to simulate a semantic disambiguation task similar to MacGregor et al. (32). The task assesses the semantic ambiguity early in a sentence, which is disambiguated later in the sentence on half of the trials. Speech inputs to the model were synthesized short sentences adapted from MacGregor et al. (32).

In the next section we describe the speech stimuli, present the generative model, and briefly describe the approximate inversion of the generative model as well as the two information theoretic measures that could be related to measurable brain activity.

#### 1. Speech stimuli

In the original design of MacGregor and colleagues, eighty sentence sets were constructed to test the subjects’ neural response to semantic ambiguity and disambiguation. Each set consists of four sentences in which two sentence MIDDLE WORDS crossed with two sentence <u>final words</u>. From the two sentence middle words, one was semantically ambiguous and from the two sentence final words one disambiguated the ambiguous middle word, and the other did not resolve the ambiguity. For example:
*The man knew that one more ACE might be enough to win <u>the tennis</u>*.
*The woman hoped that one more SPRINT might be enough to win <u>the game</u>*.
The middle word was either semantically ambiguous (“ace” can be a special serve in a tennis game, or a poker card) or not (“sprint” only has one meaning of fast running); the two ending words either resolved the ambiguity of the middle word (“tennis” resolves “ace” to mean the special serve, not the poker card) or not (“game” can refer to either poker or tennis game). We chose this set as part of input stimuli to the model, but reduced the sentences to essential components for simplicity:
*One more ACE/SPRINT wins <u>the tennis/game</u>*.
The four sentences point to a minimum of two possible contexts, i.e. the nonlinguistic backgrounds where they might be generated: all combinations can result from a “tennis game” context, and the ACE-game combination can additionally result from a “poker game” context. Importantly, in our model the context is directly related to the interpretation of the word “ace”.
To balance the number of plausible sentences for each context, we added another possible mid-sentence word “joker”, which unambiguously refers to a poker card in the model’s knowledge. We also introduced another possible sentence structure to add syntactic variability within the same contexts:
*One more ACE/SPRINT is <u>surprising/enough</u>*.
The two syntactic structures correspond to two different types of a sentence: the “win” sentences describe an event, whereas the “is” sentences describe a property of the subject.
We chose a total of two sentence sets from the original design. The other set (shortened version) is:
*That TIE/NOISE ruined the <u>game/evening</u>*.
In these sentences, the subject “tie” can either mean a piece of cloth to wear around the neck (“neckband” in the model) or equal scores in a game. The ending word “game” resolves it to the latter meaning, whereas “evening” does not disambiguate between the two meanings. Similar to set 1, we added the possibility of property-type sentences. Table 2 lists all possible sentences and their corresponding contexts within the model’s knowledge (ambiguous and resolving words are highlighted).
The input to the model consisted of acoustic spectrograms that were created using the Praat (104) speech synthesizer with British accent, male speaker 1.
In this work we are not focusing on timing or parsing aspects, rather on how information is incorporated into the inference process in an incremental manner and how the model’s estimates about a preceding word can be revised upon new evidence during speech processing. Therefore, we chose the syllable as the interface unit between continuous and symbolic representations, and fixed the length of the input to simplify the model construction (see details in Generative model). Each sentence consists of four lemma items (single words or two-word phrases), and each lemma consists of three syllables. All syllables were normalized in length by reducing the acoustic signal to 200 samples.
Specifically, in Praat, we first synthesized full words, then separated out syllables using the TextGrid function. A 6-by-200 time-frequency (TF) matrix was created for each unique syllable by averaging its spectro-temporal pattern into 6 log-spaced frequency channels (roughly spanning from 150 Hz to 5 kHz) and 200 time bins in the same fashion as in Hovsepyan et al. (26). Each sentence input to the model was then assembled by concatenating these TF matrices in the appropriate order. Since we fixed the number of syllables in each word (Ns = 3), words consisting of fewer syllables were padded with “silence” syllables, i.e. all-zero matrices. During simulation, input was provided online in that 6-by-1 vectors from the padded TF matrix representing the full sentence were presented to the model one after another, at the rate of 1000 Hz. In effect, all syllables were normalized to the same duration of 200ms. The same TF matrices were used for the construction of the generative model as speech templates (see section 2c for details).

#### 2. Generative model

The generative model goes from a nonlinguistic, abstract representation of a message defined in terms of semantic roles to a linearized linguistic sentence and its corresponding sound spectrogram. The main idea of the model is that listeners have knowledge about the world that explains how an utterance may be generated to express a message from a speaker.
In this miniature world, the modeled listener knows about a number of *contexts*, the scenarios under which a message is generated (to distinguish them from names given to representation levels in the model, we will use *italic* to refer to factors at each level; see below). Each message can either be of an “event” *type* that describes an action within the context, or of a “property” *type* that expresses a characteristic of an entity that exists in the context. *Context* and *type* are nonlinguistic representations maintained throughout the message but make contact with linguistic entities via semantics and syntax, which jointly determine an ordered sequence of lemma that then generates the acoustic signal of an utterance that evolves over time.
As in the real world, connections from context to semantics and semantics to lemma are not one-to-one, and ambiguity arises, for example, when two semantic items can be expressed as the same lemma. In this case the model can output exactly the same utterance for two different messages. When the model encounters such an ambiguous sentence during inference, it will make its best guess based on its knowledge when ambiguity is present (see Model inversion). For illustrative purposes, we only consider a minimum number of alternatives, sufficient to create ambiguity, e.g. the word “ace” only has two possible meanings in the model. Also, while the model generates a finite set of possible sentences, they are obtained in a compositional fashion; they are not spelled out explicitly anywhere in the model, and must be incrementally constructed according to the listener’s knowledge.
Specifically, the generative model (Figure 1A) is organized in three hierarchically related submodels that differ in their temporal organization, with each submodel providing empirical priors to the subordinate submodel, which then evolves in time according to its discrete or continuous dynamics for a fixed duration (as detailed below). Overall, this organization results in six hierarchically related levels of information carried by a speech utterance, from high to low (L_1_-L_6_) we refer to them as: context, semantics and syntax, lemma, syllable, acoustic, and the continuous signal represented by time-frequency (TF) patterns that stands for the speech output signal.
Each level in the model consists of one or more factors representing the quantities of interest (e.g., *context, lemma, syllable*…), illustrated as rectangles in Fig 1A. We use the term “states” or hidden states to refer to the values that a factor can take (e.g. in the model the factor *context* can be in one of four states {‘poker game’, ‘tennis game’, ‘night party’, ‘racing game’}. For a complete list of factors and their possible states of context to lemma levels see Table 1).
As an example, to generate a sentence to describe an event under a “tennis game” *context*, the model picks “tennis serve” as the agent, “tennis game” as the patient, and “win” as their relationship. When the syntactic rule indicates that the current semantic role to be expressed should be the agent, the model selects the lemma “ace”, which is then sequentially decomposed into three syllables /eis/, /silence/, /silence/. Each syllable corresponds to eight 6-by-1 spectral vectors that are deployed in time over a period of 25 ms each. The generative model therefore generates the output of continuous TF patterns as a sequence of “chunks” of 25 ms.
We next describe in detail the three submodels:

a. Discrete non-nested: context to lemma via semantic (dependency) and syntax (linearization) The context level consists of two independent factors: the *context c* and the sentence *type Ty*. Together, they determine the probability distribution of four semantic roles: the *agent s^A^*, the *relation s^R^*, the *patient s^P^*, and the *modifier s^M^*. An important assumption of the model is that states of *context, type* and semantic roles are maintained throughout the sentence as if they had memory. These semantic roles generate a sequence of lemmas in the subordinate level, whose order is determined by the *syntax*, itself determined by the sentence *type*. This generative model for the first to the n^th^ lemma is (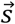 denotes the collection of all semantic factors 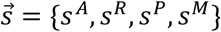:

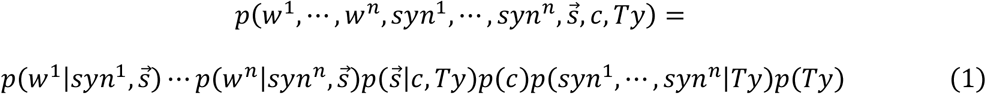 Here, p(c) is the prior distribution for the *context*. The prior probability for the sentence type p(Ty) was fixed to be equal between “property” and “event”. The terms 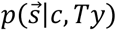 and *p*(*syn*^1^,…, *syn^n^*|*Ty*) can be further expanded as:

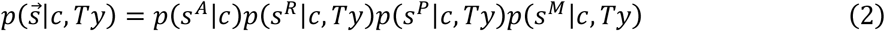

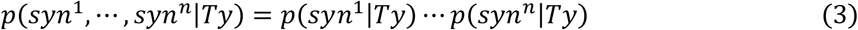 When *Ty*=‘event’, the sentence consists of an *agent*, a *patient*, a *relation* between the *agent* and the *patient*, and a null (empty) *modifier*. When *Ty*=‘property’, the sentence consists of an *agent*, a *modifier* that describes the *agent*, a *relation* that links the *agent* and the *modifier*, and a null *patient*. To translate the static context, type and semantic states into ordered lemma sequences, we constructed a minimal (linear) syntax model consistent with English grammar. We constrain all possible sentences to have four syntactic elements syn^1^-syn^4^, values are {‘attribute’, ‘subject’, ‘verb’, ‘object’, ‘adjective’}. The probability of syn^n^ is dependent solely on Ty. The syntactic element syn^i^ is active during the i^th^ epoch, and each possible value of the syntax (except ‘attribute’ that directly translates to a lemma item randomly determined within {‘one more’, ‘that’}) corresponds to one semantic factor (semantic factors in the model include subject, verb, object and adjective): Subject—*agent*; Verb—*relation*; Object—*patient*; Adjective—*modifier* Thus, sentences of the “event” type are always expressed in the form of subject-verb-object (SVO), and those of the “property” type in the form of subject-verb-adjective (SVadj). In the i^th^ lemma epoch, the model picks the current semantic factor via the value of syni and finds a lemma to express the value (state) of this semantic factor, using its internal knowledge of mapping between abstract, nonlinguistic concepts to lexical items (summarized in the form of a dictionary in Appendix I). Note that the same meaning can be expressed by more than one possible lemma, and several different meanings can result in the same lemma, causing ambiguity. The mapping from L_2_ to L_3_ can be defined separately for each lemma as follows:

- The first lemma (w^1^ the attribute) does not depend on semantics or syntax and the model would generate “one more” or “that” with equal probability (p=0.5).
- w^2^ and w^3^ are selected according to *agent* and *patient* values, respectively, which are themselves constrained by context.
- w^4^ can be either a patient or a modifier depending on Ty. Prior probabilities of context and type, as well as probabilistic mappings between levels (eq.2-4), are all defined in the form of multidimensional arrays. Detailed expressions and default values can be found in Appendix II.
b. Discrete nested: lemma to spectral Over time, factors periodically make probabilistic transitions between states (not necessarily different). Different model levels are connected in that during the generative process, discrete hidden (true) states of factors in a superordinate level (Ln) determine the initial state of one or more factors in the subordinate level (Ln+1). The Ln+1 factors then make a fixed number of state transitions. When the Ln+1 sequence is finished, L_n_ makes one state transition and initiates a new sequence at Ln+1. State transitioning of different factors within the same level occurs at the same rate. We refer to the time between two transitions within each level as one **epoch** of the level. Thus, model hierarchies are temporally organized in that lower levels evolve at higher rates and are nested within their superordinate levels. The formal definition of the discrete generative model is shown in eq.1, where the joint probability distribution of the m^th^ outcome modality (here generally denoted by *o^m^*, specified in following sections) and hidden states (generally denoted by s^n^) of the n^th^ factor up to a time point τ, is determined by the priors over hidden states at the initial epoch P(s^n, 1^), the likelihood mapping from states to outcome P(o | s) over time 1:τ, and the transition probabilities between hidden states of two consecutive time points P(s^n, t^|s^n, t-1^) up to t=τ:

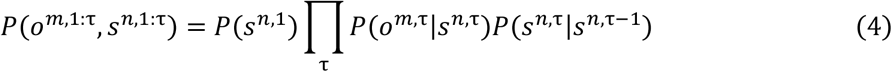 For lower discrete levels, representational units unfold linearly in time, and a sequence of subordinate units can be entirely embedded within the duration of one superordinate epoch. Therefore, the corresponding models are implemented in a uniform way: the hidden state consists of a “what” factor that indicates the value of the representation unit (e.g. the lemma ‘the tennis’), and a “where” factor that points to the location of the outcome (syllable) within the “what” state (e.g. the 2^nd^ location of ‘tennis’ generates syllable ‘/nis/’). During one epoch at each level (e.g. the entire duration of the lemma “the tennis”), the value of the “what” factor remains unchanged with its transition probabilities set to the unit matrix. The “where” factor transitions from 1 to the length of the “what” factor, which is the number of its subordinate units during one epoch (three syllables per lemma). Together, the “what” and “where” states at the lemma level generate a sequence of syllables by determining the prior for “what” and “where” states in each syllable. In the same fashion, each syllable determines the prior for each spectral vector. Thus, the syllable level goes through 8 epochs, and for each epoch the output of the syllable level corresponds to a spectral vector of dimension (1 x 6, number of frequency channels). This single vector determines the prior for the continuous submodel. Such temporal hierarchy is roughly represented in Figure 1B (downward arrows). Unlike L_1_ and L_2_ states that are maintained throughout the sentence, states of the lemma level and below are “memoryless”, in that they are generated anew by superordinate states at the beginning of each epoch. This allows us to simplify the model inversion (see next section) using a well-established framework that exploits the variational Bayes algorithm for model inversion (71). The dynamic expectation maximization (DEM) framework of Friston et al. (71) consists of two parts: hidden state estimation and action selection. In our model, the listener does not perform any overt action (the state estimates do not affect state transitioning), therefore the action selection part is omitted. Using the notation of Eq.1, parameters of the generative model are defined in the form of multidimensional arrays: Probabilistic mapping from hidden states to outcomes:

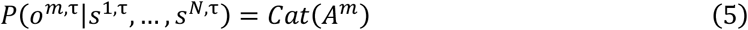 Probabilistic transition among hidden states:

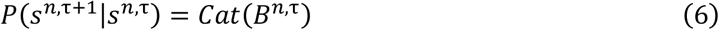 Prior beliefs about the initial hidden states:

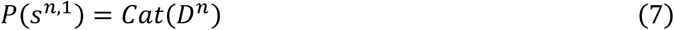 For each level we define **A, B, D** matrices according to the above description of hierarchical “what” and “where” factors:

- Probability mappings (matrix **A**) from a superordinate “what” to a subordinate “what” states are deterministic, e.g. p(sylb=‘/one/’|lemma=‘one more’, where=1)=1, and no mapping is needed for “where” states;
- Transition matrices (**B**) for “what” factors are all identity matrices, indicating that the hidden state does not change within single epochs of the superordinate level;
- Transition matrices for “where” factors are off-diagonal identity matrices, allowing transition from one position to the next;
- Initial states (**D**) for “what” factors are set by the superordinate level, and always start at position 1 for “where” factors.
c. Continuous: acoustic to output The addition of an acoustic level between the syllable and the continuous levels is based on a recent biophysically plausible model of syllable recognition, Precoss (26). In that model syllables were encoded with continuous variables and represented, as is the case here, by an ordered sequence of 8 spectral vectors (each vector having six components corresponding to six frequency channels). In the current model we only implemented the bottom level of the Precoss model (see also (28)), which deploys spectral vectors into continuous temporal patterns. Specifically, the outcome of the syllable level sets the prior over the hidden cause, a spectral vector **I** that drives the continuous model. It represents a chunk of the time-frequency pattern determined by the “what” and “where” states of the syllable level s^ω^ and s^γ^ respectively:

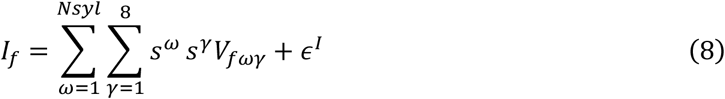

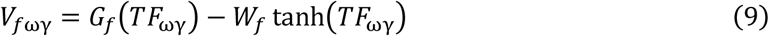 The noise terms ε^I^ is random Gaussian fluctuation. TF_ωγ_ stands for the average of the 6×200 TF matrix of syllable ω in the γ^th^ window of 25 ms. **G** and **W** are 6×6 connectivity matrices that ensure the spectral vector **I** determines a global attractor of the Hopfield network that sets the dynamics of the 6 frequency channels. Values of **G**, **W** and a scalar rate constant κ in eq. 9-10 are the same as in Precoss:

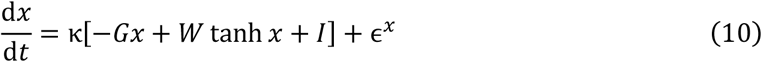 The continuous state of **x** determines the final output of the generative model **v**, which is compared to the speech input during model inversion. As **x, v** is a 6×1 vector:

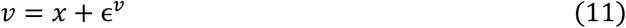 The precision of the output signal depends on the magnitude of the random fluctuations in the model (ε in eq. 8, 10, 11). During model inversion, the discrepancy between the input and the prediction of the generative model, i.e. the prediction error, are weighted by the corresponding precisions and used to update model estimates in generalized coordinates (41). We manipulated the precisions for continuous state **x** and activities of frequency channels **v** to simulate from intact (HP) to impaired (LP) periphery. The precision for top-down priors from the syllable level, Ps, was kept high for all simulations (see Table 1 for values used in different conditions). The continuous generative model and its inversion were implemented using the ADEM routine in the SPM12 software package (105), which integrates a generative process of action. Because we focus on passive listening rather than interacting with the external world, this generative process was set to identical to the generative model and without an action variable. Precisions for the generative process were the same for all simulations (Table 4).

**Table 4.** Precisions.

| Precision | Generative model: HP | Generative model: LP | Generative process |
| --- | --- | --- | --- |
| $P^x$ | $\exp(16)$ | $\exp(6), \exp(0), \exp(-4)$ | $\exp(16)$ |
| $P^v$ | $\exp(16)$ | $\exp(6), \exp(0), \exp(-4)$ | $\exp(16)$ |
| $P^l$ | $\exp(8)$ | $\exp(8)$ | $\exp(8)$ |

#### 3. Model inversion

The goal of the modeled listener is to estimate posterior probabilities of all hidden states given observed evidence p(s| o), which is the speech input to the model, here represented by TF patterns sampled at 1000 Hz. This is achieved by the inversion of the above generative model using the variational Bayesian approximation under the principle of minimizing free energy (106). Although this same computational principle is applied throughout all model hierarchies, the implementation is divided into three parts corresponding to the division of the generative model. Because the three “submodels” are hierarchically related we follow and adapt the approach proposed in (71), which shows how to invert models with hierarchically related components through Bayesian model averaging. The variational Bayes approximation for each of the three submodels is detailed below.
Overall, the scheme results in a nested estimation process (Figure 1B). For a discrete-state level L_n_, probability distributions over possible states within each factor are estimated at discrete times over multiple inference epochs. Each epoch at level L_n_ starts as the estimated L_n_ states generate predictions for initial states in the subordinate level L_n+1_, and ends after a fixed number of state transitions (epochs) at L_n+1_. State estimations for L_n_ are then updated using the discrepancy between the predicted and observed L_n+1_ states. The L_n_ factors make transitions into the next epoch immediately following the update, and the same process is repeated with the updated estimation. Different model hierarchies (from L_2_ on) are nested in that the observed L_n+1_ states are state estimations integrating information from L_n+2_ with the same alternating prediction-update paradigm, but in a faster timescale. A schematic of such a hierarchical prediction-update process is illustrated in Figure 1B.
Since levels “lemma” to the continuous acoustic output conform to the class of generative models considered in (71), we use their derived gradient descent equations and implementation. Levels “context” and “semantic and syntax” do not conform to the same class of discrete models (due to their memory component and non-nested temporal characteristics); we therefore derived the corresponding gradient descent equations based on free energy minimization for our specific model of the top two levels Equations 2-4 (see Appendix III for the derivation) and incorporated them into the general framework of DEM (71).
The variational Bayes approximation for each of the three submodels is detailed below.

a. Lemma to context For all discrete-state levels, the free energy F is generally defined as (106):

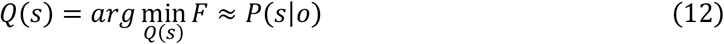

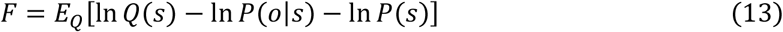 In eq. 12 and 13, Q(s) denotes the estimated posterior probability of hidden state s, P(o | s) the likelihood mapping defined in the generative model, and P(s) the prior probability of s. The variational equations to find the Q(s) that minimizes Free energy can be solved via gradient descent. We limit the number of gradient descent iterations to 16 in each update to reflect the time constraint in neuronal processes. Although context/type and semantic/syntax are modeled as two hierarchies, we assign them the same temporal scheme for the prediction-update process at the rate of lemma units, i.e. they both generate top-down predictions prior to each new lemma input, and fulfill bottom-up updates at each lemma offset. Therefore, it is convenient to define their inference process in conjunction. The posterior distribution 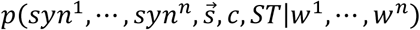 is approximated by a factorized one, *Q*(*syn*^1^)… *Q*(*syn^n^*)*Q*(*s*^1^)… *Q*(*s^n_s_^*)*Q*(*c*)*Q*(*ST*), and is parameterized as follows: 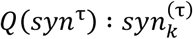, or 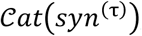, *k* = 1,…, # of possible syntactic elements, *τ* = 1,…, *n* 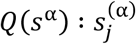, or 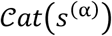, *j* = 1,…, # of possible states for semantic factor,

*α* = {*A, R, P, M*} *Q*(*c*): *c_m_*, or 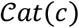, *m* = 1,…, # of possible states for context factor *Q*(*Ty*): *Ty_a_*, or 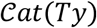, *a* = 1,…, # of possible states for sentence type Here, the model observation is the probability of the word being w^τ^ given the observed outcome o^τ^, p(w^τ^| o^τ^), which is gathered from lower-level models described in next sections. We denote p(w^τ^| o^τ^) by a vector W_i_^τ^, where τ stands for the epoch, and *i* indexes the word in the dictionary. At the beginning of the sentence, the model predicts the first lemma input, which is, by definition, just one of the two possible attributes, ‘one more’ or ‘that’.

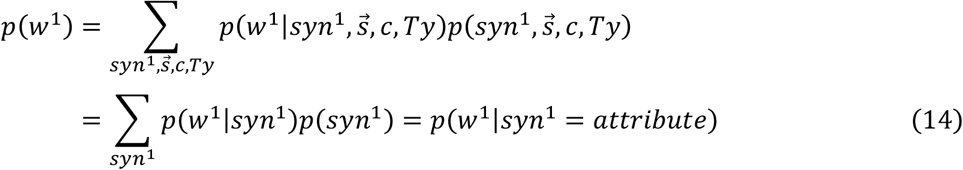 The lower levels then calculate p(w^1^ |o^1^) and provide an updated Wi^1^ that incorporates the observation made from the first lemma. This is passed to the top levels to update L_1_ and L_2_ states. Following this update, the next epoch is initiated with the prediction for w^2^. Because w^2^ does not directly depend on lemma inputs before and after itself, we can derive the following informed prediction of w^2^ from eq.2, where prior for L_1_ and L_2_ factors are replaced by their updated posterior estimates:

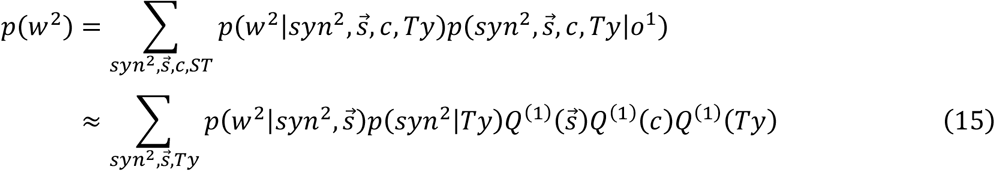 Where we used:

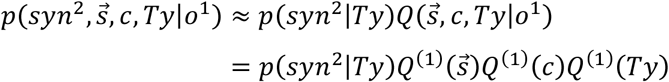 During the second epoch, the model receives input of the second lemma and updates the estimation of W_i_^2^. The updated Wi^2^ is then exploited to update L_1_ and L_2_ states, which in turn provides the prediction for w^3^. The process is repeated until the end of the sentence. The updating of L_1_ and L_2_ states, i.e. the estimation of their posterior probabilities after receiving the n^th^ lemma input relies on the minimization of the total free energy F_1,2_ of the two levels (L1, L_2_)

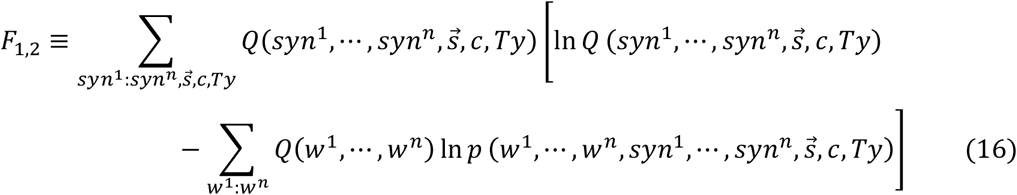 The expanded expression of F_1,2_ and derivation of the gradient descent equations can be found in Appendix III.
b. Spectral to lemma The memoryless property of lower-level (lemma and below) states implies that the observation from the previous epoch does not directly affect the prediction for the new epoch, only indirectly through the evidence accumulated at superordinate levels. The framework from Friston et al. (71) is suitable for such construction. It uses the same algorithm of free-energy (inserting eq. 5-7 to eq. 12-13) minimization for posterior estimation, but this time there is conditional independence between factors in the same level. We implemented this part of the model by adapting the variational Bayesian routine in the DEM toolbox from the SPM12 software package.
c. Continuous to spectral To enable the information exchange between the continuous and higher discrete levels that were not accounted for in (26), we implemented the inversion of the spectral-to-continuous generative model using the “mixed model” framework in (71). Essentially, the dynamics of spectral fluctuation determined by each spectral vector **I** (eq.8) is treated as a separate model of continuous trajectories, and the posterior estimation of **I** constitutes post-hoc model comparison that minimizes free energy in the continuous format. For a specific model m represented by spectral vector Im, the free energy F(t)m can be computed as (adapted from (71)):

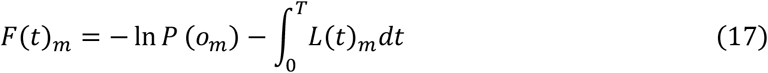

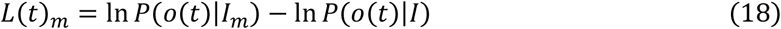 P(o_m_) indicates the likelihood for the m^th^ spectral vector (discrete). P(o(t)|Im) is the likelihood of observing the continuous input o(t) given the m^th^ **I** vector, and P(o(t)|I) is the averaged likelihood over all possible **I** vectors. In this way, the model compares the top-down prediction of I and the estimate derived from the bottom-up evidence of integrated acoustic input over 25ms. Detailed explanation of the algorithm can be found in previous studies (71, 107). The software implementation was also adapted from existing routines in the DEM toolbox of SPM12 (105).

#### Information theoretic metrics

Two metrics were derived from the belief updating process just described: the Kullback-Leibler (KL) divergence (Div), which characterizes the discrepancy between the current and previous state estimates of a factor, and entropy H that characterizes the uncertainty of the current state estimates of the factor. We denote the posterior probability of the i^th^ possible state of an arbitrary factor at time point τ as q^t^_i_. The divergence and entropy are defined as:

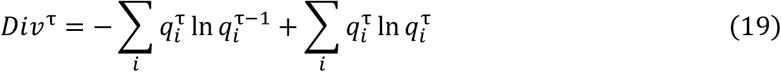

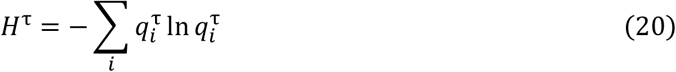

These two (non-orthogonal) metrics provide a qualitative summary of the model response that can be linked to neurophysiological signals (see Result and Discussion).

## Model guided MEG data analysis

### Next-word prediction statistics from GPT-2 model

We implemented a transformer pre-trained language model, GPT-2 (20) in Google Colab (108), to obtain word prediction statistics of the sentence stimuli. The model is trained on ~40 GB text data and generates next-word predictions given arbitrary sentence contexts. Inputs to the model were sentences taken from (32), each sentence consisting of four parts (see Table 3 for an example set): a lead-in phrase, a target word, a bridge phrase, and a resolution word. For every lead-in phrase, four variations were played by crossing two different Target words and two different Resolution words.

#### Target

either with or without semantic ambiguity (Ambiguous vs. Unambiguous).

#### Resolution

either resolves the semantic ambiguity of the Ambiguous Target, or not (Resolve vs. Unresolve).

For each set of (Target × Resolution) combination, two versions of the lead-in phrase were available. However, only one of the two lead-ins in each set was used for each subject in the MEG experiment, i.e. each set of (Target × Resolution) combination was played only once. Therefore, we averaged the GPT-2 prediction metrics for the two versions. The bridge phrase was the same within each set, regardless of other parts of the sentence.

The original speech stimuli in (32) contained sentence sets where the Target words were ambiguous between two phonetically identical but morphologically different words. These sets were removed for the GPT-2 analysis as well as for the MEG data analysis, resulting in 58 out of 80 sets.

Probability distributions of the next-word prediction of GPT-2 were obtained for two time points to calculate the prediction entropy and surprisal, respectively:

1. After Target, i.e. the input to GPT-2 is [lead in] + [target] We use the entropy H of this prediction as a proxy for the (semantic) ambiguity of the target word, with the hypothesis that if a word has multiple meanings, different meanings will predict different next words with similar probabilities, resulting in a flatter distribution compared to the prediction from its unambiguous counterpart. H is calculated as follows, where *i* indexes all words in the dictionary:

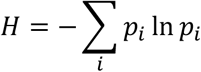
2. Before Resolution, i.e. the input to GPT-2 is [lead in] + [target] + [bridge] We calculate the surprisal S for each resolution word from the prediction probability as follows, where r is the index for the resolution word in the dictionary:

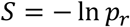 This surprisal is equivalent to the KL divergence of the posterior distribution after the resolution word, because the distribution has collapsed to p=1 for the received word and 0 elsewhere.

### MEG sensor space analysis

The MEEG module in SPM12 (105) was used for the MEG data preprocessing. Statistical analysis and plotting of the preprocessed results were performed with the Fieldtrip Toolbox (109). We first performed the identical preprocessing as MacGregor et al. (32) on head-adjusted raw MEG responses to the 58 selected sentence sets for all 16 subjects. Briefly, raw recordings were first bandpass filtered between 0.1 and 40 Hz, then epoched at the offsets of each keyword (Target or Resolution). After baseline correction and the rejection of bad trials, combined gradiometer (RMS of each of the 102 gradiometer pairs) responses were cropped into shorter time windows (−0.2~0.8s for the Target offset, −0.5~1s for the Resolution offset) and averaged across trials for each subject. For averaging, trials were split in the following way that allow for statistical tests for both the GPT-2 prediction metrics and the linguistic metrics of interest, i.e. semantic ambiguity at the Target offset and resolution at the Resolution offset:

1. Target Sentences were split into two groups: 1. The GPT-2 entropy for the Ambiguous word was larger than the entropy for the Unambiguous word (Amb1, Uam1), and 2. The GPT-2 entropy for the Ambiguous word was smaller than for the Unambiguous word (Amb2, Uam2).
2. Resolution Sentences containing the Resolve words were split into two groups: 1. The GPT surprisal of the Resolve word following the Ambiguous target was larger than the Resolve word following the Unambiguous target (Res_Amb1, Res_Uam1), and 2. The GPT surprisal of the Resolve word following the Ambiguous target was smaller than following the Unambiguous target (Res_Amb2, Res_Uam2).

To assess the effects of linguistic and GPT-2 metrics on the combined gradiometer data, we constructed the following four contrasts:

1. [Amb1 + Amb2] vs. [Uam1 + Uam2]: effect of semantic ambiguity.
2. [Amb1 + Uam2] vs. [Amb2 + Uam1]: effect of GPT-2 prediction entropy.
3. [Res_Amb1 + Res_Amb2] vs. [Res_Uam1 + Res_Uam2]: effect of preceding ambiguity.
4. [Res_Amb1 + Res_Uam2] vs. [Res_Uam1 + Res_Amb2]: effect of GPT-2 prediction surprisal.

To test for differences between the two conditions within each contrast, we first took the average of the two averages in each condition within individual subjects, e.g. (Amb1 + Amb2)/2 for the ambiguous condition in contrast 1. This yields one sensor × time response per condition and per subject. We then performed a paired t-test across subjects for each sensor and time point, resulting in a 2D parametric map of the test statistic. Clusters of sensors with ps <0.05 were identified on this map, each including at least 2 neighboring sensors. The statistical significance of each cluster was evaluated by comparing the maximum t-statistic of the cluster to a null distribution generated by randomly permuting the condition labels within each subject (5000 times across all 16 subjects). The cluster-level p-value (pc) was the proportion of the t statistic in the permutation distribution larger than the maximum t statistic of the selected cluster. None of the clusters identified by the t-test survived the permutation test, therefore we report the five clusters with the highest t-statistics for the positive effect in each contrast. We also computed Cohen’s d (110) from the grand average over time and across subjects of all the 102 combined gradiometer channel to evaluate the effect size of each contrast at single gradiometer pairs.

## Acknowledgements

We thank B. Bickel, S. van Ommen, D. Poeppel for critical feedback, NCCR TTF Data Science for support on the GPT-2 model, and E. Holmes for advice on the SPM software. This work was funded by Swiss National Science Foundation (grant number 320030B_182855) and NCCR Evolving Language, Swiss National Science Foundation Agreement #51NF40_180888.

## Data and code availability

Custom MATLAB code and simulation data will be made available upon request.

## Supporting Figures

**S1 Fig.**
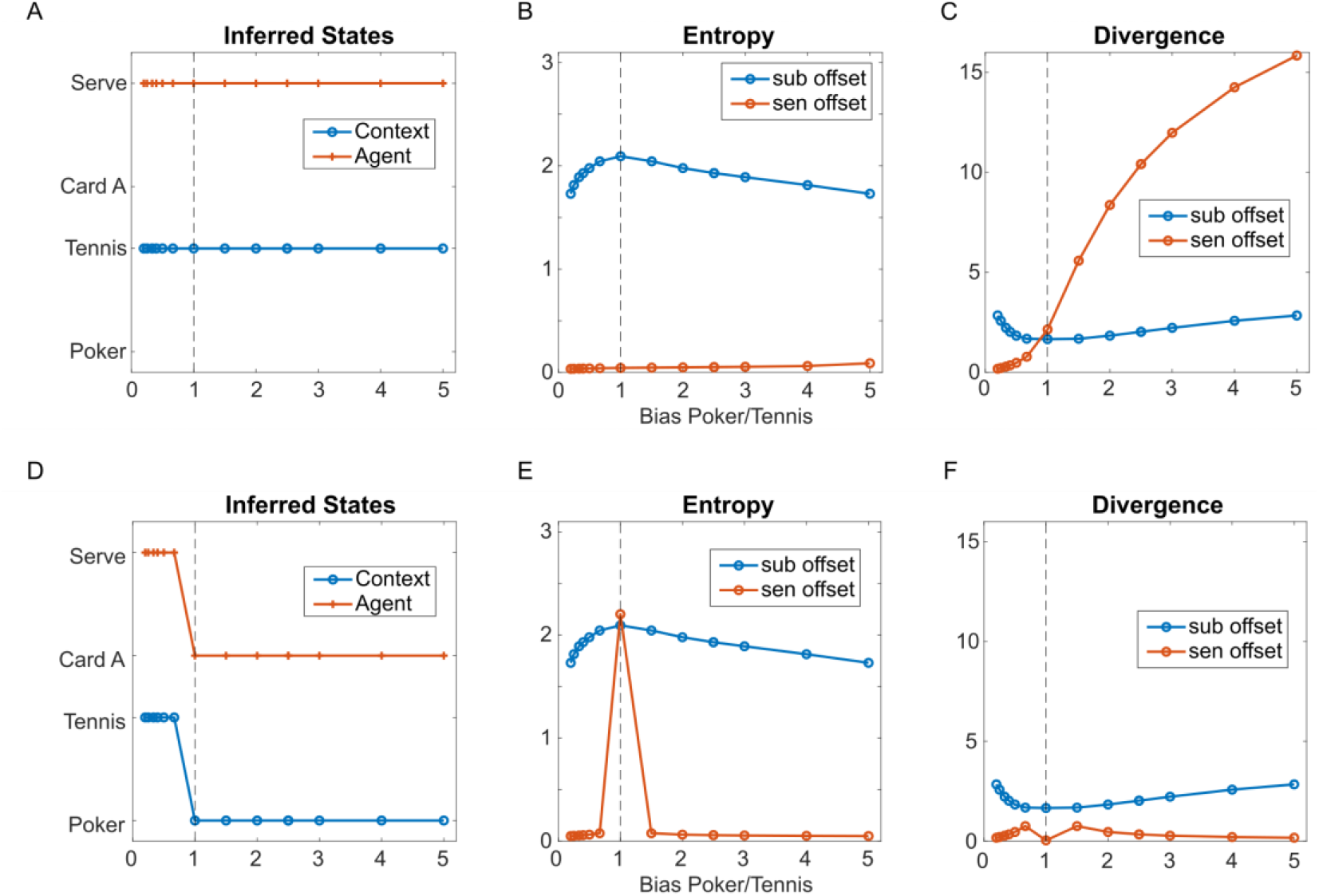
Effect of contextual bias ratio on the inference process. **A-C: metrics derived from the sentence “One more ace wins the tennis” as function of contextual bias between “poker game” and “tennis game”.** A bias of x implies that the prior probability ratio (the total probability is always normalized to 1) for context was set to [x 1 1 1] for all 4 possible contexts {‘poker game’, ‘tennis game’, ‘night party’, ‘racing game’} for x>=1, and [1 1/x 1 1] for x<1 to balance the influence of the two irrelevant contexts. **D-F: same metrics derived from sentence “One more ace wins the game”. A.** Inferred states for the *context* (blue) and the *agent* (red) do not change with contextual bias, i.e. the model always resolved to the correct states. **B.** Sum of entropy across *context, agent* and *patient* at the subject word (“ace”) offset and the sentence offset. At the offset of “ace” (blue), the entropy is maximum at bias=1 and symmetric on both sides. At sentence offset (red), the entropy is overall lower than at the offset of “ace” and monotonically increases with a small slope, reflecting that the model was more certain about the state estimations at this point, but keeps a small possibility towards the poker game that increases with the bias towards the poker context. **C.** At the sentence offset, the divergence monotonically increases with bias towards poker reflecting the increasing difference between the expected context (poker) and the actual one (tennis). **D.** Inferred states for context and agent at the end of sentence B as a function of bias. For bias<1 (preference for ‘tennis’context), the inferred context is “tennis (game)” and inferred agent is “serve”. For bias>=1, the result corresponds to a preference for the “poker” context. **E.** Sum of entropy. For both time points, the entropy is at maximum when bias=1. Both curves are symmetrical by bias=1. The blue curve is the same as in B because the sentence input up to this point was the same. **F.** Sum of divergence across the same three factors at two critical time points. At the offset of “ace”, the divergence reached its minimum at bias=1 as a result of the uniform distribution over “poker” and “tennis” states, which is the least different from the previous time point. At the sentence offset, the stronger the bias (farther from 1), the smaller the difference between before and after hearing the final word. However, a notch is seen at bias=1 due to the uncertainty (S1 Fig E).

**S2 Fig.**
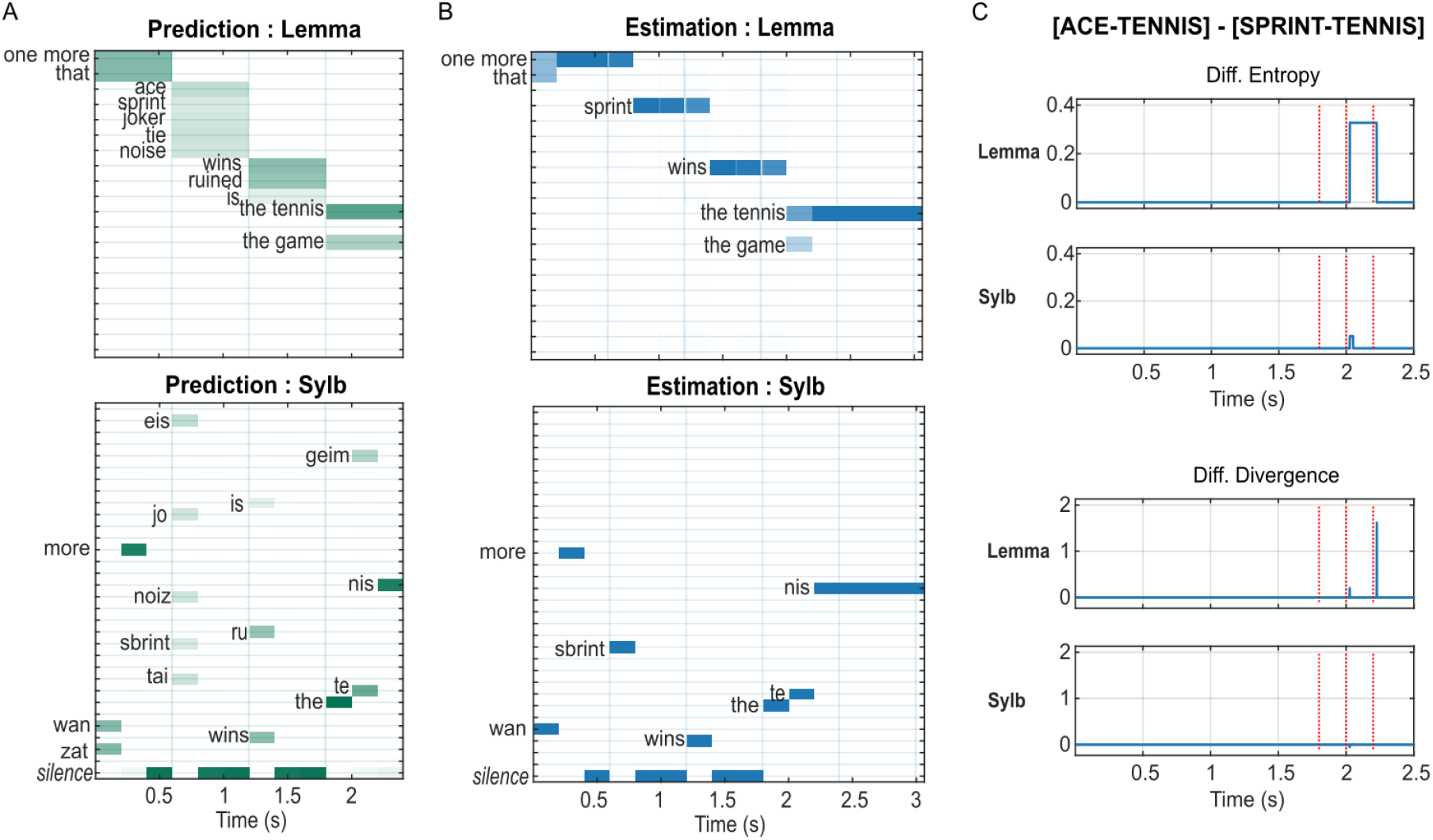
Message passing in the processing of the same word in different sentences. Figure specifications are the same as Fig 3. **A. Semantic-to-lemma and lemma-to-syllable predictions in response to sentence “one more sprint wins the tennis”.** The second lemma “sprint” influences the prediction for the final lemma as well as the corresponding syllables as compared to Fig 3A. **B. Estimation of posterior probabilities for lemma and syllable states for the sentence [SRPINT-tennis].** Similar to Fig 3B, the model instantly recognizes each syllable (lower panel). **C. Upper panels: entropy derived from sentence [ACE-TENNIS] minus sentence [SPRINT-TENNIS] for the lemma and the syllable levels for the entire sentence.** Vertical dotted lines mark the onset of each syllable of the final lemma. Entropies for both the lemma and the syllable level was higher for [ACE-TENNIS] after the onset of the second syllable, reflecting a greater complexity (three possible states compared to two in the sentence [SPRINT-TENNIS]) of the prediction of the final lemma. **Lower panels: the difference between the divergence in response to the two sentences.** A positive difference at the onset of the third syllable (the offset of the second syllable) indicates that the input “the tennis” is less expected in the sentence [ACE-TENNIS] due to the prior preference for the poker context, compared to in the sentence [SPRINT-TENNIS] where the context was already resolved to “poker game” after hearing “sprint”.

**S3 Fig.**
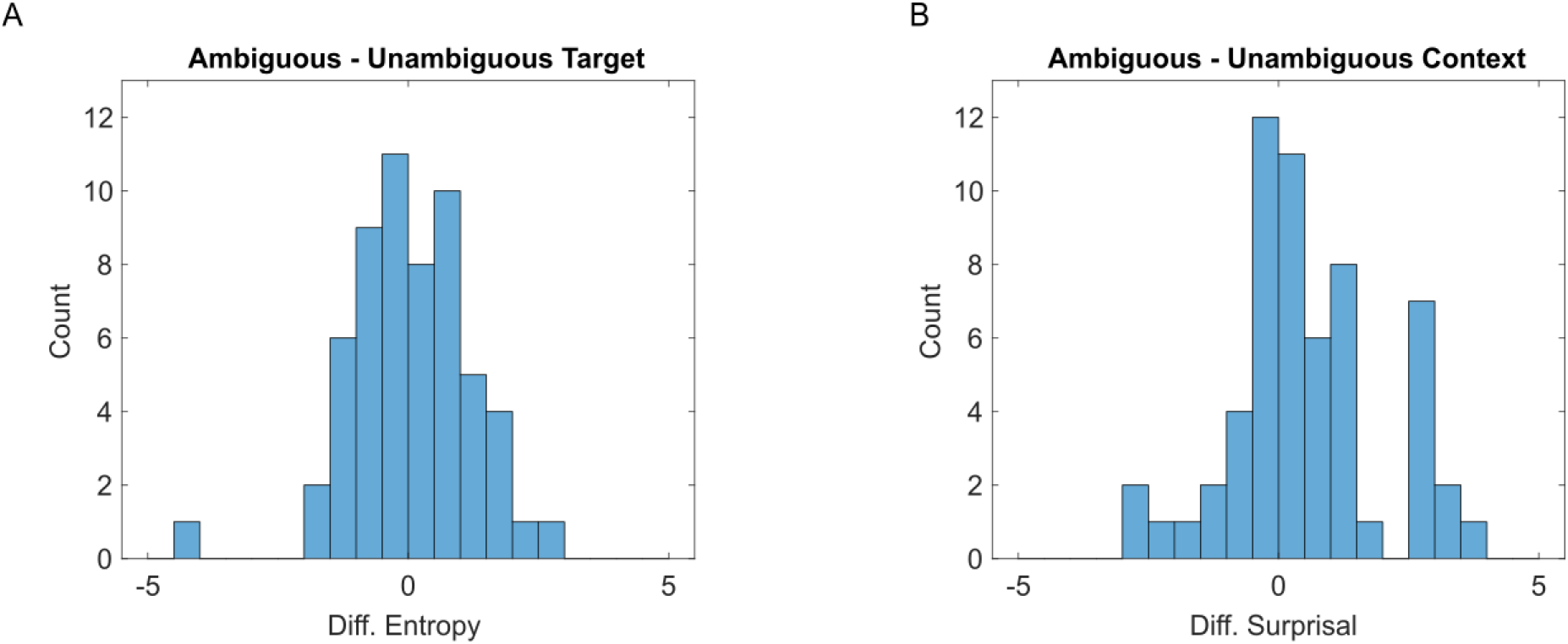
**A.** Distribution for the difference of GPT-2 prediction entropy calculated from ambiguous vs. unambiguous Target words. Only the 58 selected sentences were included. **B.** Distribution for the difference of GPT-2 prediction surprisal calculated from the same Resolution words following ambiguous vs. unambiguous Target.

**S4 Fig.**
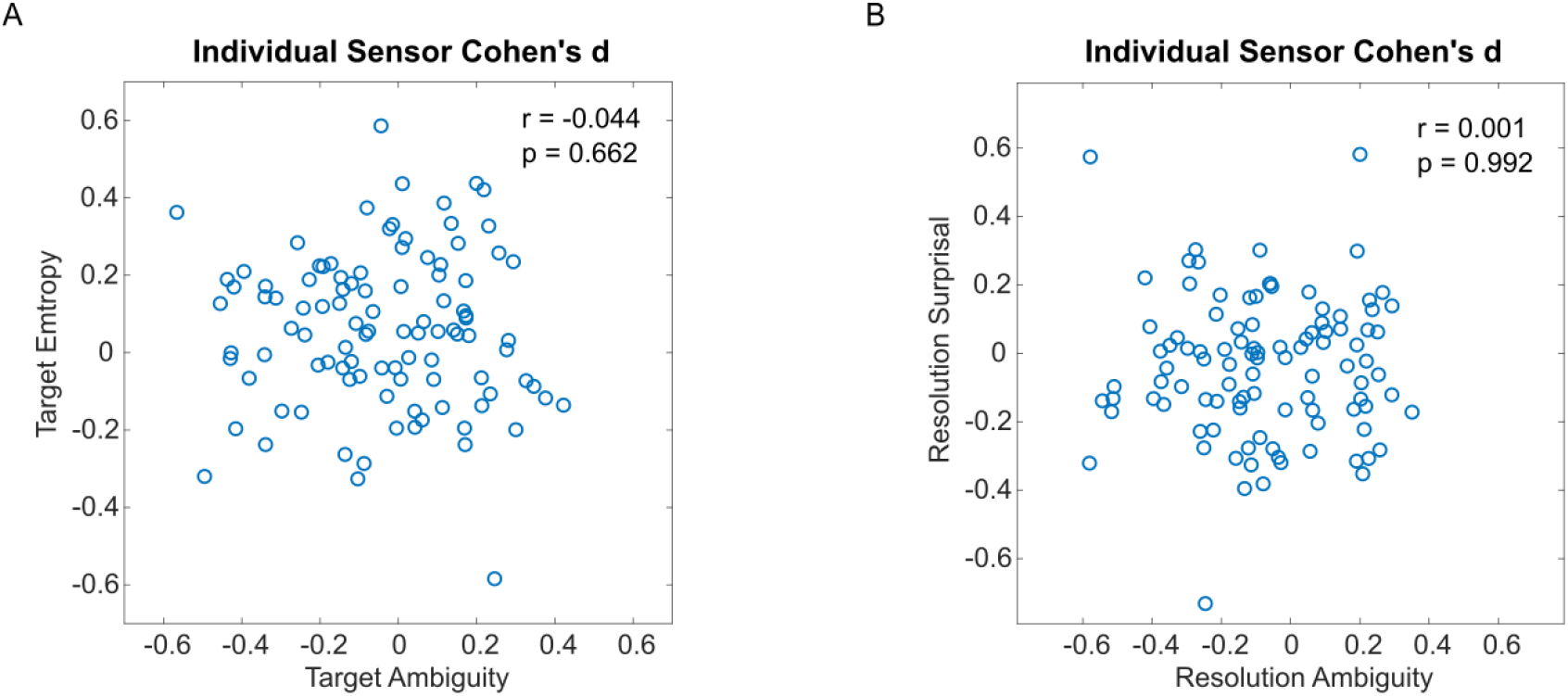
Comparison of effect sizes between semantic and GPT-2 prediction metrics. A. Cohen’s d computed from the effect of semantic ambiguity (x-axis) and the effect of GPT-2 prediction entropy (y-axis) at Target offset for each of the 102 combined gradiometers. B. Cohen’s d for the effect of preceding ambiguity (x-axis) vs. GPT-2 prediction surprisal (y-axis) at Resolution offset for each combined gradiometer.

**S5 Fig.**
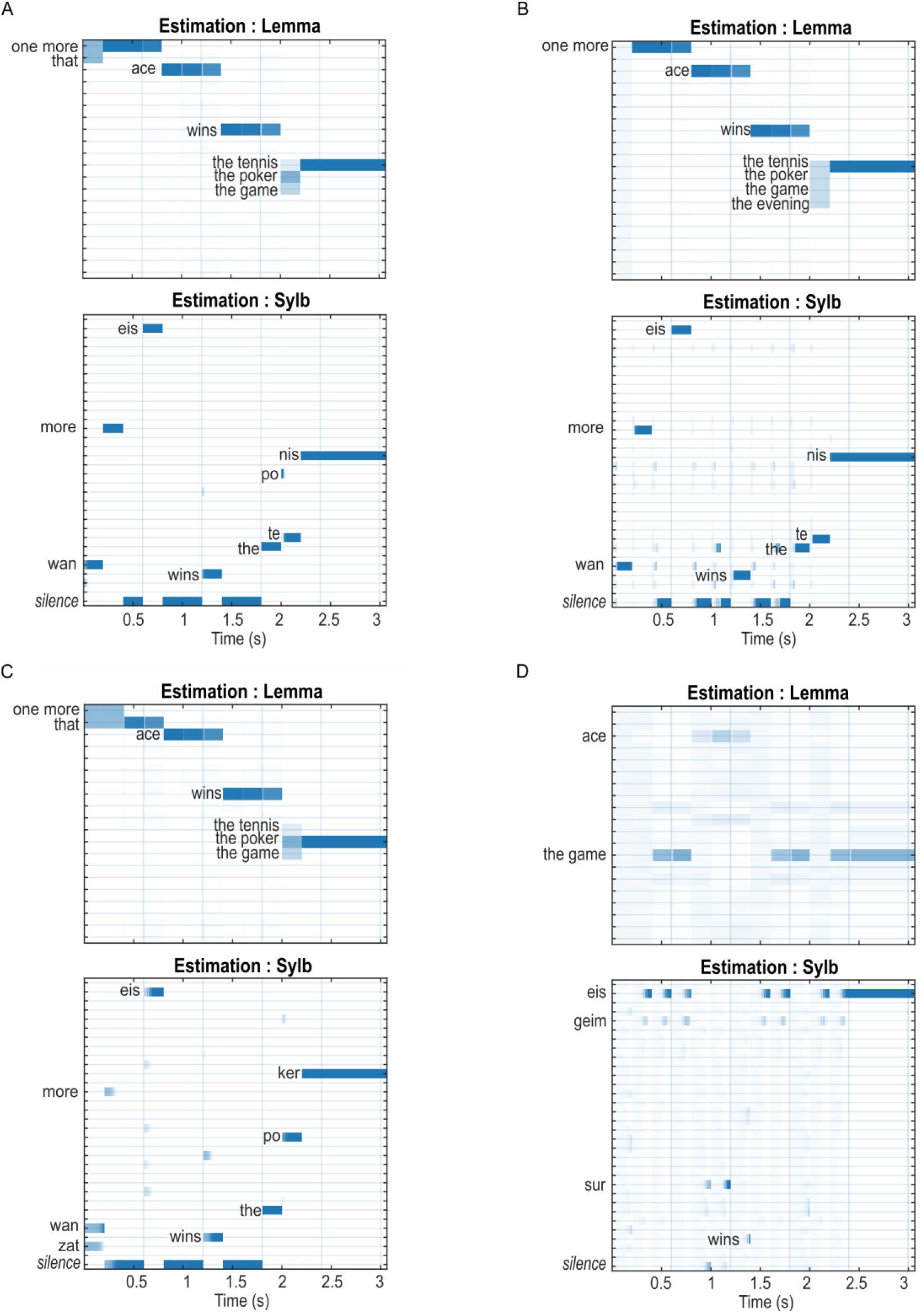
A, B: Inference of lemma and syntax states at moderately high precision (exp(6)) with (A) or without (B) informative top-down predictions. The posterior estimates are very similar to the intact condition (Fig 4B and 5A, respectively) in that the model quickly converged onto the correct states after each update. However, longer delays to convergence can be observed at the syllable level with prediction, and both lemma and syllable levels without prediction, compared to their intact counterparts. **C, D: Inference of lemma and syntax states at extremely low precision (exp(−4)) with (C) or without (D) informative top-down predictions.** The posterior estimates with informative prediction are qualitatively the same as the low-precision condition in Fig 6A but with longer delays before convergence. Without any top-down prediction, the model completely fails at the syllable level, hence cannot make accurate estimates for higher levels.

