## Appendix for "A deep hierarchy of predictions enables assignment of semantic roles in online speech comprehension"

**Appendix I. Model parameters for lemma generation**

1. Possible states and prior belief (D matrix) for each factor
a. Context level

**Context** = {‘poker game’, ‘tennis game’, ‘night party’, ‘racing game’}, Nc = 4

D{1} = [1.5 1 1 1]

**Type** = {‘event’, ‘property’}, NTy = 2

D{2} = [1 1]

b. Semantic

**Agent** = {‘card A’, ‘serve’, ‘run’, ‘card J’, ‘neckband’, ‘score’, ‘buzz’}, Na = 7

**Relation** = {‘win’, ‘ruin’, ‘be’}, Nr = 3

**Patient** = {‘tennis’, ‘poker’, ‘game’, ‘evening’, ‘null’}, Np = 5

**Modifier** = {‘sufficient’, ‘unexpected’, ‘not pretty’, ‘not fair’, ‘high volume’, ‘high freq’, ‘null’}, Nm = 7

Prior belief for semantic and syntax factors are calculated by multiplying the higher-level priors with probability mapping matrices defined in the next section.

c. Syntax

For syntax, we treat each epoch differently as if there is one factor syntax{i} for each epoch.

**Syntax**{1-4} = {‘attribute’, ‘subject’, ‘verb’, ‘object’, ‘adjective’}, Nsyn = 5

d. Lemma

**Lemma** = {‘one more’, …, ‘sharp’}, Nw = 20. The matching between lemma and syntax + semantic is defined in Appendix I. Multiple meanings are arranged assuming meaning 1 is the major meaning, meaning 2 & 3 are less likely.

2. Probabilistic mapping for the generative model

In all mapping matrices, the first dimension represents the outcome factor, the 2^nd^ and further dimensions represent states of the higher level. All matrices are normalized so that the first dimension add to 1.

a. Context to semantic

For the mapping from context c and sentence type Ty to a semantic factor s (s={ a’, ‘r’, ‘p’, ‘m’}), the model is defined by a 3-D matrix L{s}(Ns, Nc, NTy). Indices refer to the state of the factor, e.g. L{agent}(1, 1, 2) = p(agent=’card A’|context=’poker’, type=’property’).

**Context = ‘poker game’ = 1:**

L{agent}(‘card A’, 1, 1) = L{agent}(‘card A’, 1, 2) = 0.6, L{agent}(‘card J’, 1, 1) = L{agent}(‘card J’, 1, 2) = 0.4

L{relation}(‘win’, 1, 1) = 1, L{relation}(‘be’, 1, 2) = 1

L{patient}(‘poker’, 1, 1) = 1, L{patient}(‘null’, 1, 2) = 1

L{modifier}(‘null’, 1, 1) = 1, L{modifier}(‘sufficient’, 1, 2) = L{modifier}(‘unexpected’, 1, 2) = 0.5

**Context = ‘tennis game’ = 2:**

L{agent}(‘serve’, 2, 1) = L{agent}(‘serve’, 2, 2) = 0.6, L{agent}(‘run, 2, 1) = L{agent}(‘run’, 2, 2) = 0.4

L{relation}(‘win’, 2, 1) = 1, L{relation}(‘be’, 2, 2) = 1

L{patient}(‘tennis’, 2, 1) = 1, L{patient}(‘null’, 2, 2) = 1

L{modifier}(‘null’, 2, 1) = 1, L{modifier}(‘sufficient’, 2, 2) = L{modifier}(‘unexpected’, 2, 2) = 0.5

**Context = ‘night party’ = 3:**

L{agent}(‘neckband’, 3, 1) = L{agent}(‘neckband’, 3, 2) = 0.6, L{agent}(‘buzz’, 3, 1) = L{agent}(‘buzz’, 3, 2) = 0.4

L{relation}(‘ruin’, 3, 1) = 1, L{relation}(‘be’, 3, 2) = 1

L{patient}(‘evening’, 3, 1) = 1, L{patient}(‘null’, 3, 2) = 1

L{modifier}(‘null’, 3, 1) = 1

L{modifier}(‘not pretty’, 3, 2) = L{modifier}(‘high volume’, 3, 2) = L{modifier}(‘high freq’, 3, 2) = 1/3

**Context = ‘racing game’ = 4:**

L{agent}(‘score’, 4, 1) = L{agent}(‘score’, 4, 2) = 0.6, L{agent}(‘buzz’, 4, 1) = L{agent}(‘buzz’, 4, 2) = 0.4

L{relation}(‘ruin’, 4, 1) = 1, L{relation}(‘be’, 4, 2) = 1

L{patient}(‘game’, 4, 1) = 1, L{patient}(‘null’, 4, 2) = 1

L{modifier}(‘null’, 4, 1) = 1

L{modifier}(‘not fair’, 4, 2) = L{modifier}(‘high volume’, 4, 2) = L{modifier}(‘high freq’, 4, 2) = 1/3

b. Sentence type to syntax

For syntax, we define the model Z for each epoch τ separately. Each Z{τ} is a Nsyn x NTy matrix, i.e. 5x2

Z{1}(:, 1) = Z{1}(:, 2) = [1 0 0 0 0]. The first epoch is always ‘attribute’

Z{2}(:, 1) = Z{2}(:, 2) = [0 1 0 0 0]. The second epoch is always a subject

Z{3}(:, 1) = Z{3}(:, 2) = [0 0 1 0 0]. The third epoch is always a verb

Z{4}(:, 1) = [0 0 0 1 0], Z{4}(:, 2) = [0 0 0 0 1]. The fourth epoch could be either an object (sentence type=’event’) or an adjective (Ty=’property’).

Now we can calculate priors for semantic and syntax (matrix multiplication is only demonstrative, not concerning matrix transposing in practice).

Semantic: D{3-6} = L{1-4}*D{1}*D{2}

Syntax: D{7}(:, τ) = Z{τ}*D{2}

c. Semantic to lemma

Because we have a one-to-one correspondence between syntax and semantic, we define the model A with Nsyn independent matrices, each mapping from the corresponding semantic factor to word, therefore Nw x Ns.

Special case for A{1} because it does not need semantic information. Therefore A{1} is a Nw x 1 matrix. A{1}(one more) = A{1}(that) = 0.5

A{2}(index, agent) = 1, where index is calculated by finding the dictionary entry for the corresponding agent. E.g. the 3^rd^ agent state, ‘card J’, translates to ‘joker’ in the dictionary, which is at the 5^th^ entry. Therefore A{2}(5, 3) = 1. The rest of semantic-lemma mappings are defined in the same fashion.

When one semantic value corresponds to multiple lemma entries, e.g. the patient ‘tennis’ (1^st^ patient) can be translated into either ‘the tennis’ (11^th^ lemma entry), or ‘the game’ (13^th^), we give higher probability to the “meaning 1” mapping. A{4}(11, 1) = 0.8, A{4}(13, 1) = 0.2.

**Appendix II. The mapping between lemma and semantic in the model’s mental lexicon**

|  | lemma | Meaning 1 | Meaning 2 | Meaning 3 |
| --- | --- | --- | --- | --- |
| 1 | 'one more' | 'extra' | [] | [] |
| 2 | 'that' | 'that' | [] | [] |
| 3 | 'ace' | 'card A' | 'serve' | [] |
| 4 | 'sprint' | 'run' | [] | [] |
| 5 | 'joker' | 'card J' | [] | [] |
| 6 | 'tie' | 'neckband' | 'score' | [] |
| 7 | 'noise' | 'buzz' | [] | [] |
| 8 | 'wins' | 'win' | [] | [] |
| 9 | 'ruined' | 'ruin' | [] | [] |
| 10 | 'is' | 'be' | [] | [] |
| 11 | 'the tennis' | 'tennis' | [] | [] |
| 12 | 'the poker' | 'poker' | [] | [] |
| 13 | 'the game' | 'game' | 'tennis' | 'poker' |
| 14 | 'the evening' | 'evening' | [] | [] |
| 15 | 'enough' | 'sufficient' | [] | [] |
| 16 | 'surprising' | 'unexpected' | [] | [] |
| 17 | 'ugly' | 'not pretty' | [] | [] |
| 18 | 'unfair' | 'not fair' | [] | [] |
| 19 | 'loud' | 'high volume' | [] | [] |
| 20 | 'sharp' | 'high freq' | [] | [] |

**Appendix III. Full expression of free energy and gradient descent algorithm for the top-level model (L_1_ and L_2_)**

The expression of free energy (eq.16 in the main text) can be parameterized with respect to the posterior estimates of L_1_ and L_2_ factors

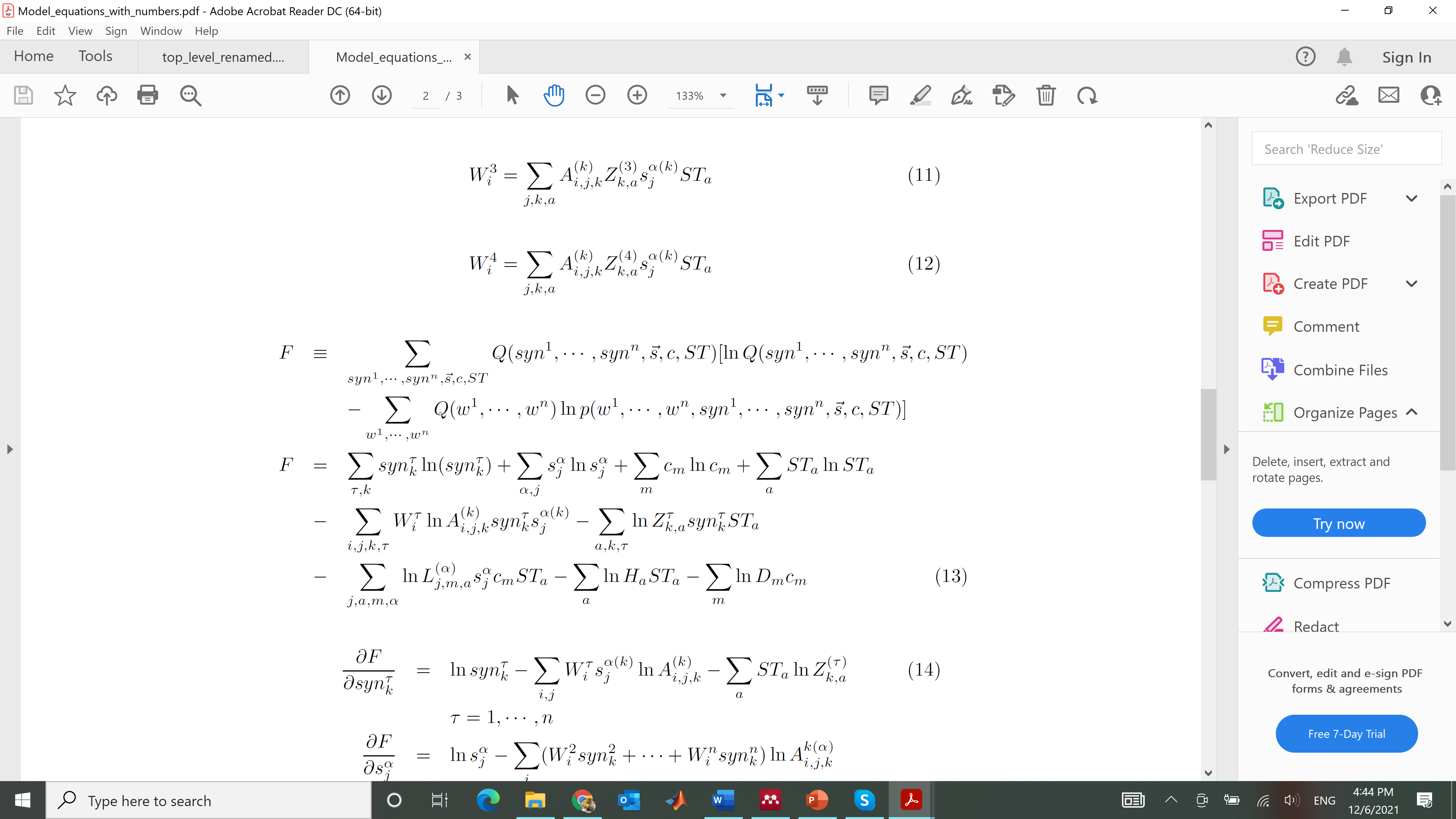

We can then derive partial derivatives of F with respect to Q:

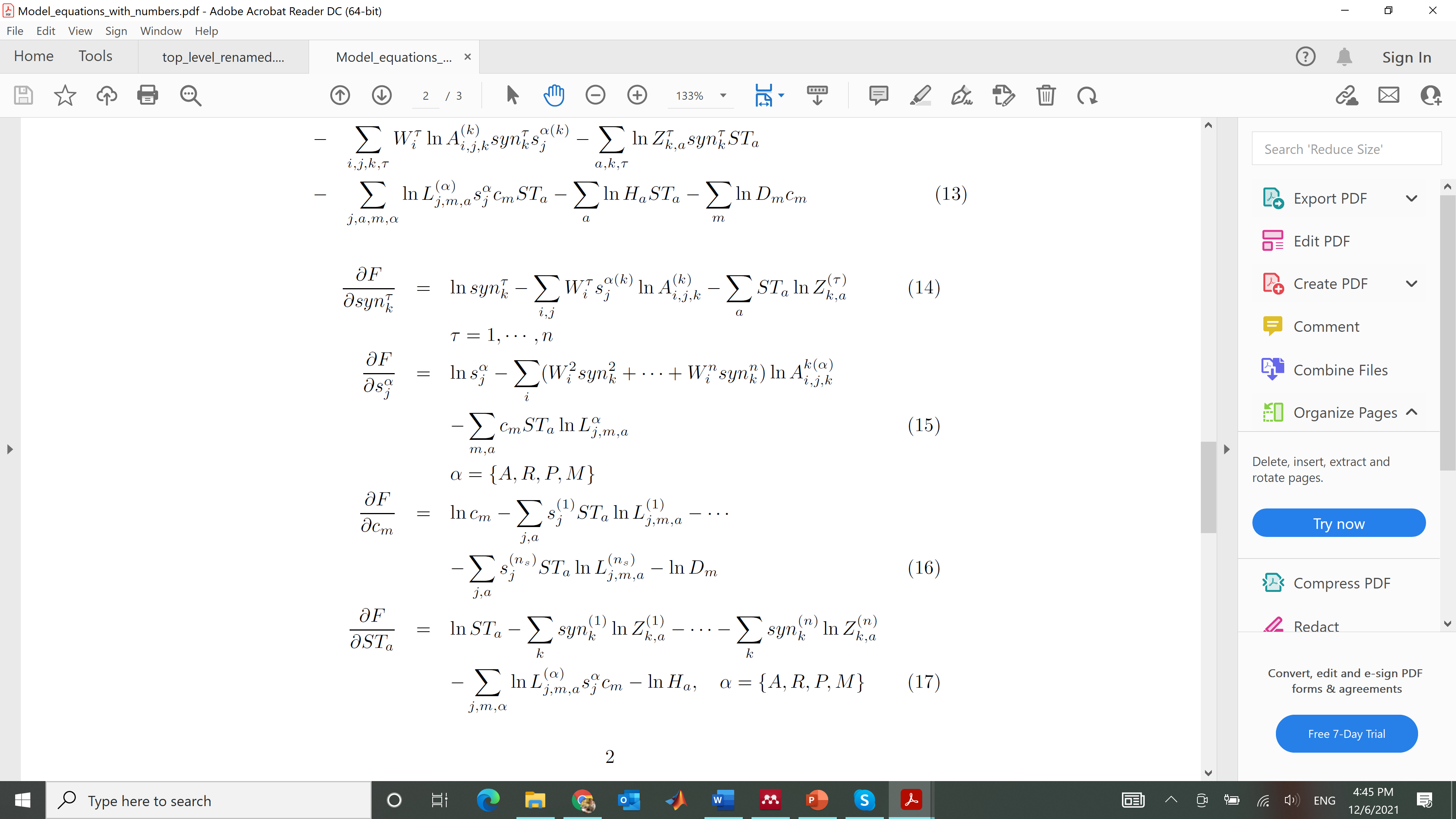

To solve the above equations, we follow Friston et al. (2017) and define an auxiliary variable *v* for the estimation of each factor *x*. Let *x* ≡ σ(*v*), where σ() denotes the softmax function. We can then solve *v* and *x* using gradient descent:

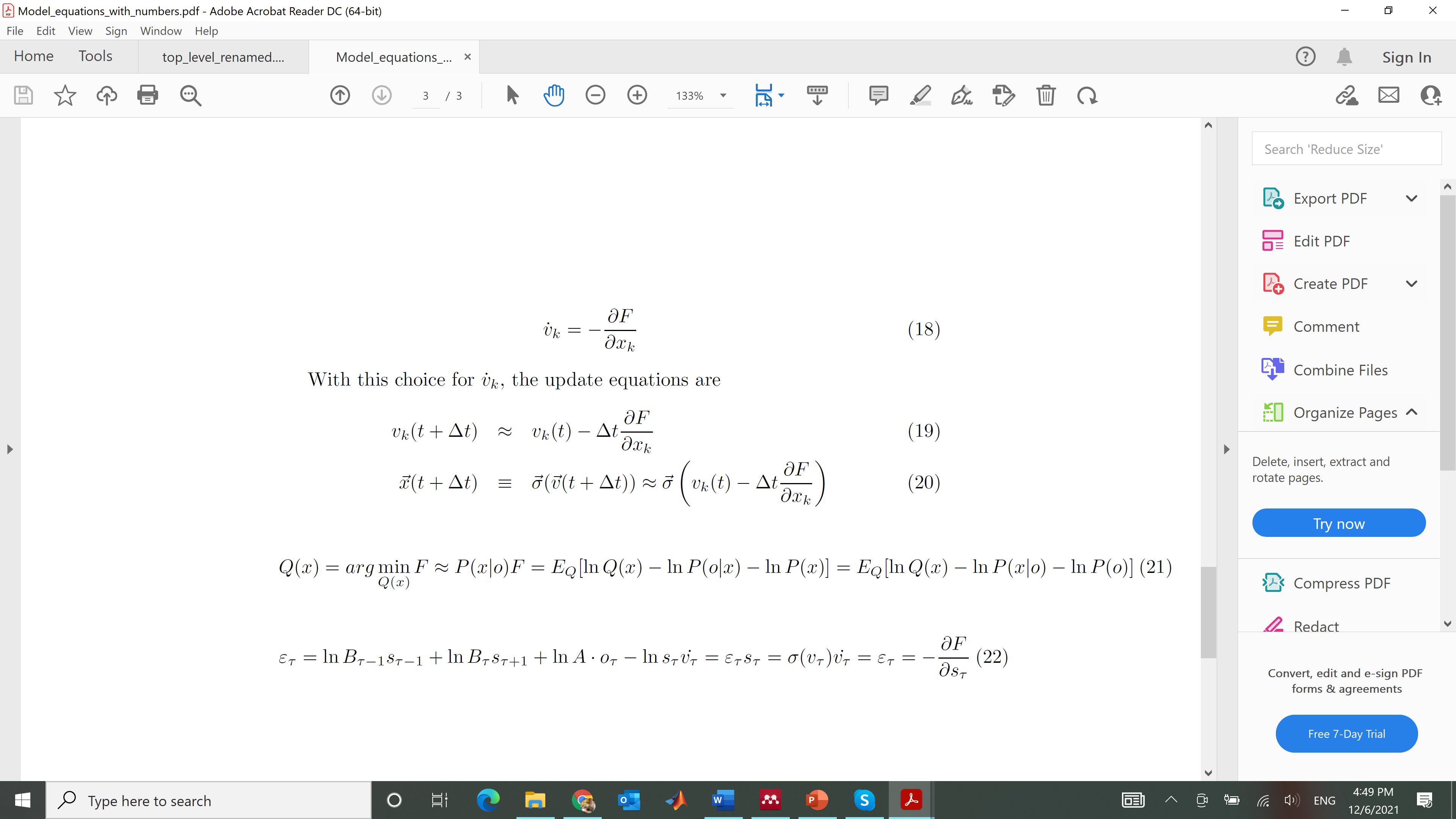
